# Interaction between BID and VDAC1 is required for mitochondrial demise and cell death in neurons

**DOI:** 10.1101/2021.09.14.460262

**Authors:** Sina Oppermann, Barbara Mertins, Lilja Meissner, Cornelius Krasel, Georgios Psakis, Philipp Reiß, Amalia M. Dolga, Nikolaus Plesnila, Moritz Bünemann, Lars-Oliver Essen, Carsten Culmsee

**Author notes:** Corresponding authors: Prof. Dr. Carsten Culmsee, Institute for Pharmacology and Clinical Pharmacy, Faculty of Pharmacy, University of Marburg, Karl-von-Frisch-Str. 2, 35032 Marburg, Germany; Dr. Sina Oppermann, Hopp Children’s Cancer Center Heidelberg (KiTZ), Heidelberg, Germany and Clinical Cooperation Unit Pediatric Oncology, German Cancer Research Center (DKFZ) and German Cancer Consortium (DKTK), Heidelberg, Im Neuenheimer Feld 280, 69120 Heidelberg, Germany.

## Abstract

Mitochondrial damage is a key feature of regulated cell death in neurons. In particular, mitochondrial outer membrane permeabilization (MOMP) has been proposed as a starting point for mitochondrial demise upon cellular stress. Potential mechanisms for MOMP presented in the literature include membrane pore formation by Bcl2-family proteins such as BID and BAX, oligomerization of voltage-dependent anion channels (VDACs) and hetero-oligomer formation of these proteins. In our study, we demonstrate a direct interaction between the voltage-dependent anion channel VDAC1 and the pro-apoptotic protein BID in dying neurons both *in vitro* and *in vivo*. Binding of BID to VDAC1 affects anion conductance through VDAC1 and is associated with glutamate-induced cell death in cultured neurons and ischemic brain injury. In cultured neurons, reducing VDAC1 expression significantly attenuates BID-induced hallmarks of mitochondrial damage such as mitochondrial fission, declined mitochondrial respiration, increased ROS production, and mitochondrial membrane potential breakdown. Our data highlight a critical role for VDAC1 as a mitochondrial receptor for activated BID, thereby serving as a key decision point between life and death in neurons.

**One Sentence Summary:** VDAC1 interacts with BID to mediate mitochondrial membrane permeabilization and neuronal cell death.

---

Regulated cell death (RCD) is a key feature of progressive neuronal loss in neurodegenerative diseases, stroke, traumatic brain injury and most other forms of chronic or acute brain damage. The mechanisms underlying programmed neuronal death include, among others, oxidative dysregulation, intracellular Ca^2+^overload and DNA damage, and finally result in loss of mitochondrial membrane potential (ΔΨ_m_) and mitochondrial disintegration, a process believed to be the “point of no return” in the cell’s commitment to die [1, 2]. Breakdown of mitochondrial integrity and function is associated with impaired oxidative phosphorylation, diminished energy supply, increased production of reactive oxygen species (ROS), and the release of mitochondrial proteins such as cytochrome c and apoptosis-inducing factor (AIF), which accelerate the final steps of programmed neuronal cell death [1, 3–6]. Mitochondrial outer membrane permeabilization (MOMP) is mediated by the formation of protein-permeable pores by pro-apoptotic members of the Bcl-2 protein family, such as BAX, BAK, BID, or BIM [7–9]. In addition, mitochondrial pathways of regulated cell death have been linked with voltage-dependent anion channels (VDAC) located in the outer mitochondrial membrane [10–13]. Three mammalian isoforms of VDAC have been identified (VDAC1, VDAC2, and VDAC3). VDAC1 is the most abundant isoform and is believed to be the principal channel-forming protein information of the mitochondrial permeability transition pore (mPTP) [14, 15]. More recently, VDAC1 has been linked to mitochondrial death pathways in response to oxidative damage in a model of glutamate-induced oxytosis in neuronal HT22 cells [16]. These findings suggested a key role for VDAC1 for both, mitochondrial permeability transition and pro-apoptotic MOMP formation. The latter may serve as a starting point for mitochondrial damage as a key decision point in programmed cell death in neurons and can be accelerated by interaction of VDAC1 with pro-apoptotic BAX or BID [17–21]. In our previous work, we have demonstrated a key role for BID in model systems of neuronal cell death including models of oxytosis and ferroptosis in HT22 cells, oxygen glucose deprivation and glutamate excitotoxicity in primary neurons in vitro, and in model systems of cerebral ischemia in vivo [22–25]. In the current study we investigated whether the pro-apoptotic bcl-2 family member BID physically interacts with VDAC1, thereby causing mitochondrial damage and neuronal cell death both *in vitro* and *in vivo*.

## Methods

### Cell culture and induction of apoptosis

HT-22 cells were cultured in Dulbeccos’s modified Eagle medium (DMEM, Invitrogen, Karlsruhe, Germany) supplemented with 10 % heat-inactivated fetal calf serum, 100 U/ml penicillin, 100 µg/ml streptomycin and 2 mM glutamine (all from PAA Laboratories GmbH, Germany). Primary mouse embryonic cortical neurons were cultured in neurobasal medium with 2% (v/v) B27 supplementary, 2 mM glutamine and 100 U/ml penicillin/streptomycin (Invitrogen). For induction of apoptosis, growth medium was replaced by medium containing glutamate and cells were analysed between 4 h and 20 h after treatment.

### Transient focal cerebral ischemia and tissue preparation

Male C57BL/6 mice (n=6) (body weight, 18–22 g; Charles River Laboratories, Sulzfeld, Germany) were subjected to 60 min transient middle cerebral artery occlusion (MCAo). Surgery was performed in isoflurane/N_2_O anesthesia (1.5% isoflurane, 68.5% N_2_O, 30% O_2_). Tissue homogenates for immunoprecipitation and western blot analysis were generated from 4 mm sections of the ipsi- and contralateral parietal cortex (2 mm +/- bregma) respectively at 2 h, 6 h and 24 h after the onset of ischemia. All procedures described are in accordance with local laws and were approved by the animal protection committee of the Government of Upper Bavaria.

### Immunoprecipitation studies

Immunoprecipitation of BID, VDAC1 and Flag-VDAC1 was performed by pull-down of the proteins from total protein lysates of HT-22 cells exposed to glutamate at the indicated time points 0 h to 16 h, from primary cortical neurons exposed to glutamate (25 µM 0 h, 4 h and 22 h) and from tissue homogenates of mice subjected to MCAo, according to the manufacturer’s protocol (Invitrogen) with the following variations. Magnetic Protein A-Dynabeads were prepared for each condition for effective binding of the respective antibodies: anti-BID (7.5 µg, Cell Signaling), anti-VDAC1 (N18) (7.5 to 10 µg, Santa Cruz, Biotechnology) or anti-Flag (7.5 μg, Sigma-Aldrich). To detect protein-interaction partners of BID and VDAC1, 30 µl of the eluate was analyzed by electrophoresis followed by western blot analysis. 30-150 μg of total protein lysate per treatment condition was used as a control in the 12.5% SDS-PAGE.

### VDAC1 gene silencing

For siRNA transfections Lipofectamine RNAiMax (Invitrogen, Karlsruhe, Germany) was used. VDAC1 siRNA or non-functional scrambled RNA was separately dissolved in Optimem I (Invitrogen, Karlsruhe, Germany). After 10 min of equilibration at RT, each siRNA solution was combined with the respective volume of the Lipofectamine RNAiMax solution and mixed gently. For reverse transfection the transfection mixture was filled in the cell culture wells/dishes and allowed to form siRNA liposome complexes for further 20 min. at RT. After addition of antibiotic-free cell suspension, a final concentration of 20 nM to 80 nM RNA and 2 µl/ml Lipofectamine was reached. Controls were treated with 100 µl/ml Opti-MEM I only and vehicle controls with 2µl/ml Lipofectamine. For siRNA sequences see extended experimental procedures.

### Analysis of cell viability of HT-22 cells

For morphological analysis of cell viability, transmission light microscopy of living HT-22 neurons was performed using an Axiovert 200 microscope (Carl Zeiss, Jena, Germany) equipped with a Lumenera Infinity 2 digital camera. Quantification of cell viability in HT-22 cells was performed either in standard 96-well plates or standard 24-well plates by MTT (3-(4,5-dimethylthiazol-2-yl)-2,5-diphenyltetrazolium bromide, Sigma-Aldrich) reduction. Absorbance was determined at 570 nm (FluoStar OPTIMA, BMG Labtech, Offenburg, Germany). Real-time detection of cellular viability was conducted by cellular impedance measurements using the xCELLigence system Real-Time Cell Analyzer RTCA-MP (Roche Diagnostics, Penzberg, Germany) [26]. Apoptotic cell death was detected by annexin-V/propidium iodide staining and subsequent flow cytometry analysis using the Guava Easy Cyte 6-2 L system (Merck Millipore, Schwalbach, Germany). Annexin-V-FITC emission was detected with the green filter at 525/530 nm, fluorescence of PI with the red filter at 690/650 nm [27].

### Analysis of energy production and oxygen consumption rate in HT-22

For analysis of total ATP levels, VDAC1 siRNA transfected and non-transfected HT-22 neurons were seeded in white 96-well plates (Greiner, Frickenhausen, Germany) for luminescence measurements. Twenty-four hours after seeding, cells were treated with 5 mM glutamate. Cellular ATP levels were detected 20 h after glutamate exposure using the ViaLight™ Plus-Kit (Lonza, Verviers, Belgium) following the manufacturer’s instructions. The values were given as relative values in % of control cells. For OCR measurements HT-22 cells were seeded in XF96-well cell culture microplates (Seahorse Bioscience) at 10,000 cells/well in standard medium and incubated at 37 °C and 5% CO_2_ for ~20 h. Before measurements, growth medium was washed and replaced with ~180 μl of assay medium (with 4.5g/L glucose as the sugar source, 2 mM glutamine, 1 mM pyruvate, pH 7.35) and cells were incubated at 37 °C for 60 min. Following the recording of three baseline measurements, Oligomycin at a final concentration of 3 µM, FCCP at a concentration of 0.4 µM and Rotenone/Antimycin A at a concentration of 1 µM were injected in the ports A-C respectively. Three measurements were performed after the addition of each compound.

### Analysis of Lipid peroxidation

For detection of cellular lipid peroxidation, cells were stained with 2 µM BODIPY 581/591 C11 (Invitrogen) in standard medium and incubated at 37 °C for 60 min. Afterwards, cells were harvested, washed and resuspended in PBS and measured by flow cytometry (FACS) using the Guava Easy Cyte 6-2 L system (Merck Millipore, Schwalbach, Germany). Fluorescence was excited at 488nm and BODIPY emission was recorded at 530 and 585 nm. Increases in green fluorescence indicated formation of lipid peroxides.

### Analysis of mitochondrial membrane potential

Loss of mitochondrial membrane potential ΔΨ_m_ was determined by using the MitoPT™ TMRE kit (Immunochemistry Technologies, Hamburg, Germany). Cells were incubated for 20 min at 37 C with TMRE (tetramethylrhodamine ethyl ester) after glutamate treatment. Cells were collected, washed with PBS and resuspended in assay buffer. As a positive control for a complete loss of ΔΨ_m_, the protonophore carbonyl cyanide m-chlorophenylhydrazone (CCCP, 50 µM) was applied to intact cells. Flow cytometry was performed using an excitation at 488 nm and an emission and 680 nm.

### Immunostaining and confocal laser scanning fluorescence microscopy

For fluorescence analysis, HT-22 cells were seeded in Ibidi µ-slide 8-well plates (Ibidi, Munich, Germany) at a density of 1.6 – 2.0 × 10^4^ cells per well. Twenty-four hours after seeding, cells were transfected with FLAG-VDAC1 in presence and absence of BI6c9 (10 µM) or treated with glutamate. Mitochondria were stained with MitoTracker DeepRed according to the manufacturer’s protocol (Invitrogen) or visualized by co-transfection with a mitochondria-targeted GFP construct (MGFP). End-point microscopic pictures were taken after fixation of the cells with 4% (v/v) paraformaldehyde (PFA) 17-24 h after treatment, followed by FLAG-VDAC1 immunostaining using a specific monoclonal FLAG M2 antibody (anti-FLAG M2, Sigma-Aldrich) followed by the secondary DyLight 649 anti-mouse antibody (Merck Millipore, Darmstadt, Germany) or the Alexa Fluor 488 antibody (green). The specificity of the respective Flag-VDAC1 immune reactivity was controlled by emission of the primary antibody in parallel staining of negative controls. Images were acquired using a confocal laser scanning microscope (Leica SP5, Wetzlar, Germany). Light was collected through a 63 × 1.4 NA, oil immersion objective. MGFP was excited at 488 nm and emissions were detected using 505-530 band pass filter (green), DyLight-stained FLAG-VDAC1 was detected by excitation at 620 nm band pass filter and emission by using a 690 nm long pass filter (red). For quantification of mitochondrial morphology at least 500 cells per condition were counted blind towards the treatment conditions by to investigators in at least 3-4 independent experiments. For digital imaging the software LSM Image Browser 4.2.0 (Carl Zeiss, Jena, Germany) was used.

For characterization of mitochondrial morphology, three categories of mitochondrial morphology were defined: category I shows elongated tubulin-like mitochondria, which are equally distributed throughout the cytosol; category II mitochondria are defined as intermediate and round mitochondria yet without apoptotic features of the cell; category III mitochondria are characterized as strongly fragmented organelles, accompanied by an apoptotic phenotype of the cell, indicated by nuclear condensation and peri-nuclear arrangement of the mitochondrial fragments.

### Pull down of recombinant proteins

For investigation of protein-protein interactions, recombinant full length His_6_-BID (His-rBID) and caspase 8-cleaved BID (His-rtBID) were incubated separately with recombinant mVDAC1 (rVDAC1) for 1-2 h at 4°C. His-rBID without incubation with mVDAC1 and sample containing only mVDAC1 were used as negative controls. All protein samples were separately incubated with nickel-nitrilotriacetic acid-(NiNTA-) agarose and incubated for 1 h at 4°C. The protein-Ni-NTA mixture was loaded on a Poly-Prep^®^ Chromatography column (BioRad Laboratories, Munich) and washed with buffer (10 mM Tris/HCl pH 8.0, 100 mM NaCl, 0.05 % (v/v) LDAO) 2-3 times to remove unbound protein amounts. His-rBID and His-rtBID together with their binding partners were bound to Ni-NTA and eluted with buffer containing 200 mM imidazole (10 mM Tris/HCl pH 8.0, 100 mM NaCl, 0.05 % (v/v) LDAO, 200 mM imidazole). The washing fractions (W) and elution fractions (E) for each sample were collected and acetone precipitated following the Pierce manufacturer protocol for acetone precipitation. To determine protein-protein interaction the protein samples were analyzed by 12.5% SDS-PAGE followed by Coomassie staining or Western blot.

### Thermophoresis measurements of mVDAC1 and Bid/tBid

mVDAC1 was labeled using Alexa Fluor® 532 C5-maleimide (Invitrogen). 1 mM TCEP (AppliChem) were added to 100 µM protein solution and incubated for 30 min at 4°C. 1 mM label was subsequently added and samples were incubated over night at 4°C and 350 rpm in a Thermomixer (Eppendorf). Excess label was removed by using a Sephadex G25 column (GE Healthcare Bio-Science AB, Uppsala, Schweden). Protein concentration and label efficiency were measured at 280 nm and 532 nm with a Nanodrop ND-1000 (peqLab). Microscale thermophoresis (MST) measurements were performed using the Monolith NT.115 instrument (NANO TEMPER). mVDAC1 concentration was kept constant at 50 nM and Bid / tBid were added to the solution ranging from 15 nM to 500 µM. Protein solution was incubated in the dark for 20 min at RT and centrifuged (25402 g, 5 min) to remove aggregated and/or precipitated species. Standard capillaries (NANO TEMPER) were filled with 8-10 µL of the protein solution and MST curves were recorded. K_D_ values were calculated using the NT-Analysis software. Fluorescently labeled OmpG was used to rule out unspecific interactions of Bid/tBid with β-barrels.

### Single channel conductance recordings of mVDAC1 ion channels

Single channel conductance recordings were performed using the BLM technique. Once recombinant VDAC1 was added beside the BLM and a stable single channel was inserted, Bid/ tBid were added to the *cis* side. For each measurement the differences from the baseline to the stable high conductance state (S0) and the low conductance states (S1, S2, S2’) were determined with the software. Each data point of the *U*/*I*-curve was taken in account to determine the conductivity *via* the slope derived by linear regression.

### Statistical analyses

All experiments were independently repeated three to five times with n=3-8 per condition. All data are given as means ± standard deviation (s.d.). Statistical multiple comparisons were performed by analysis of variance (ANOVA) followed by Scheffé’s *post hoc* test. Calculations were performed with the Winstat standard statistical software package (R. Fitch Software, Bad Krozingen, Germany).

## Results

### Role of VDAC1 in oxidative cell death and mitochondrial damage

First, we analyzed the role of VDAC1 in mediating the hallmark features of oxidative cell death in HT22 cells, a mouse hippocampal cell line. In this model system of glutamate-induced cell death, BID mediates mitochondrial damage and cell death upon exposure to millimolar concentrations of glutamate [23, 24, 28]. Two different siRNAs significantly reduced the expression of VDAC1 in HT22 cells measured at both the mRNA and protein levels (Figures 1A, 1B and S1A, S1B). These siRNA sequences are specific for VDAC1, as the expression of VDAC2 was not affected by VDAC1 gene silencing (Figure S1C). To investigate the effect of reduced VDAC1 expression in HT22 cells, we analyzed the morphology and viability of cells following glutamate-induced toxicity. Treating cells with a scrambled (non-silencing) siRNA failed to protect the neurons from cell death (Figure 1C); in contrast, cells that were treated with VDAC1 siRNA retained their spindle-shaped morphology (Figures 1C and S1D) and were rescued from glutamate-induced toxicity, measured using fluorescence-activated cell sorting (FACS) analysis of annexin V/propidium iodide (AV/PI)-stained HT22 cells (Figures 1D and S1E-S1F) and using the MTT assay (Figures 1E and S1G). Moreover, real-time cell analysis of cellular impedance revealed that treating cells with VDAC1 siRNA confers long-lasting protection from glutamate-induced cell death (Figures 1F and S1H).

**Figure 1.**
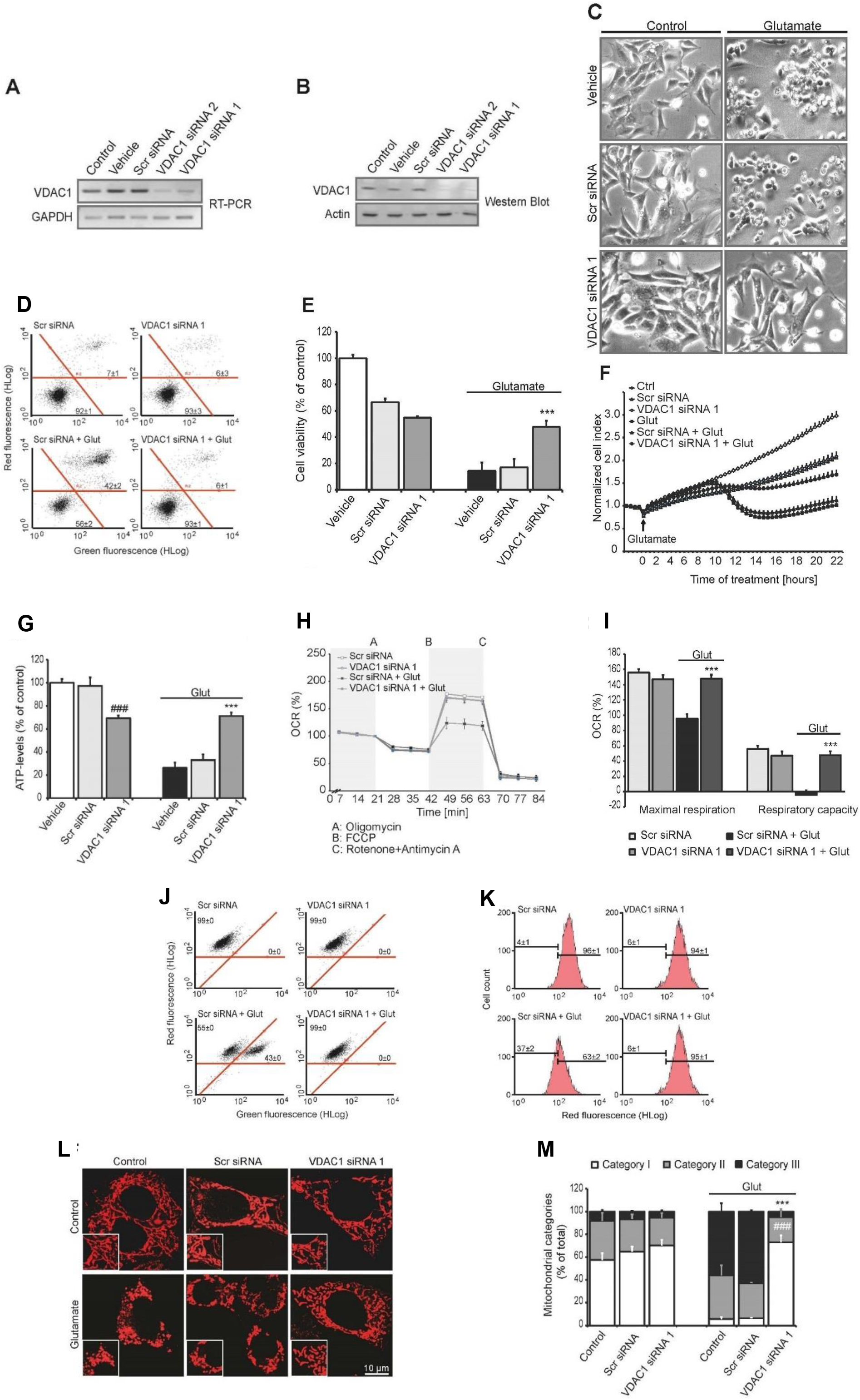
VDAC1 siRNA protects HT22 cells from glutamate-induced toxicity. **(A, B)** RT-PCR analysis of VDAC1 mRNA **(A)** and western blot analysis of VDAC1 protein levels **(B)** in HT22 cells 48 h after transfection with vehicle, a scrambled non-silencing siRNA (*scr* siRNA), VDAC1 siRNA 1, or VDAC1 siRNA 2. **(C)** Example images of HT22 cells (10x 0.25-NA objective) transfected as indicated prior to or after 16 h of glutamate (3 mM) stimulation. **(D)** FACS analysis of annexin V (green fluorescence)/propidium iodide (red fluorescence) ‒stained HT22 cells depicts dead cells (AV^+^/PI^+^) in the upper-right quadrant and healthy cells (AV^-^/PI^-^) in the lower-left quadrant. Glut:stimulation with 5 mM glutamate for 17.5 h. **(E)** Cell viability was measured using the MTT assay. Where indicated, 3 mM glutamate was applied for 16 h. ***, *p*<0.001 versus glutamate-treated vehicle and glutamate-treated *scr* siRNA (ANOVA, Scheffe’s test). **(F)** Real-time measurements of cellular impedance were performed using the xCELLigence system. **(G)** ATP luminescence measurements of vehicle- and siRNA-transfected HT22 cells. **(H)** Mitochondrial respiration in HT22 cells was measured as oxygen consumption rate (OCR). **(I)** Mitochondrial maximum respiration and respiratory capacity were calculated using the OCR data and are presented as a percentage. ^###^ and ***, *p*<0.001 versus the corresponding vehicle and *scr* siRNA-treated cells (ANOVA, Scheffe’s test). **(J)** BODIPY-FACS analysis of lipid peroxidation. **(K)** TMRE-FACS measurements of mitochondrial membrane potential (ΔΨ_m_) in HT22 cells 17 h after onset of treatment with 4 mM glutamate. **(L)** Analysis of mitochondrial morphology in HT22 cells treated as indicated HT22 in the presence or absence of glutamate (5 mM, 17 h). Mitochondria were visualized using the Mitotracker DeepRed stain. **(M)** Quantification of mitochondrial morphology. Category I indicates elongated mitochondria, category II indicates an intermediate morphology, and category III indicates fragmented mitochondria. ^###^ and ***, *p*<0.001 versus the corresponding glutamate-treated control and *scr* siRNA groups, respectively (ANOVA, Scheffe’s test). All summary data are presented as the mean ± standard deviation.

Next, we examined the effect of deleting VDAC1 on several additional hallmark features of programmed cell death by measuring ATP levels, mitochondrial oxygen consumption rate (OCR), and ROS formation. Consistent with previous studies [8, 29], treating cells with VDAC1 siRNA - but not vehicle or the scrambled, non-silencing siRNA - led to slightly reduced ATP levels (Figures 1G and S2C). However, cells transfected with VDAC1 siRNA were significantly protected against the glutamate-induced reduction of ATP levels compared to glutamate treated vehicle and non-silencing siRNA transfected control cells (Figures 1G and S2C). Similar results were obtained when we used OCR to measure mitochondrial metabolic function. Specifically, glutamate stimulation significantly reduced both maximum mitochondrial respiration and mitochondrial respiratory capacity in control-treated cells, whereas mitochondrial respiration was not reduced upon glutamate stimulation in cells transfected with VDAC1 siRNA (Figures 1H, 1I and S2D).We also found that reducing VDAC1 expression protected cells from increased ROS levels, reflected by reduced lipid peroxides 18 h after onset of the glutamate stimulation (Figures 1J and S2A, S2B), suggesting that VDAC1 plays a role in glutamate-induced irreversible mitochondrial damage and accelerated ROS formation.

Furthermore, FACS analysis of TMRE (tetramethylrhodamine, ethylester)-stained cells revealed that the transfection of cells with VDAC1 siRNA prevented ΔΨ_m_ breakdown (Figures 1K and S2G, S2H), which is a major feature of both glutamate-induced and tBID-induced toxicity; ΔΨ_m_ breakdown is also associated with mitochondrial fission and programmed cell death in neurons [30, 31]. Therefore, we examined whether knocking down VDAC1 expression preserves mitochondrial morphology and function in glutamate-stimulated cells. Consistent with our previous findings, glutamate stimulation triggered detrimental mitochondrial fission in HT22 cells (Figures 1L, 1M and S2E, S2F) [24, 30]; however, transfecting cells with VDAC1 siRNA prevented the glutamate-induced shift from healthy mitochondrial morphology (category I) to the peri-nuclear accumulation of the fragmented organelles (category III) (Figures 1L, 1M and S2E, S2F). Taken together, these data strongly suggest that VDAC1 plays a key role in intrinsic death pathways at the mitochondrial level during glutamate-induced cell death.

### The tBID-induced mitochondrial death pathway requires VDAC1

Reducing VDAC1 expression protected neurons from glutamate toxicity to a similar extent as silencing BID expression [23, 32] or pharmacologically blocking BID [28, 30, 33, 34]; therefore, we hypothesized that a direct interaction between VDAC1 and BID may underlie the BID-mediated effects on mitochondrial integrity and function. We examined the extent to which BID and VDAC1 are involved in intrinsic cell death. HT22 cells were transfected with a tBID-encoding plasmid (ptBID), and cell death was measured at several time points in the presence or absence of VDAC1. Overexpression of tBID induced a characteristic change in the morphology of HT22 cells; in contrast, VDAC1-silenced cells retained their normal morphology (Figures 2A and S3A). FACS analysis confirmed that VDAC1 deficiency protected cells from tBID-induced toxicity (Figures 2B and S3B, S3C). In addition, we found that VDAC1 is a necessary target for tBID-mediated ΔΨ_m_ breakdown (Figures 2C and S3D-E). TMRE staining was significantly reduced in tBID-expressing HT22 cells, indicating that expressing tBID leads to the breakdown of mitochondrial membrane permeability; this effect was prevented by VDAC1 gene silencing (Figures 2C and S3D-E). The finding that tBID alone caused neither a breakdown of ΔΨ_m_ nor cell death in VDAC1-depleted cells suggests that glutamate-induced toxicity in cultured neurons requires a direct interaction between tBID and VDAC1.

**Figure 2.**
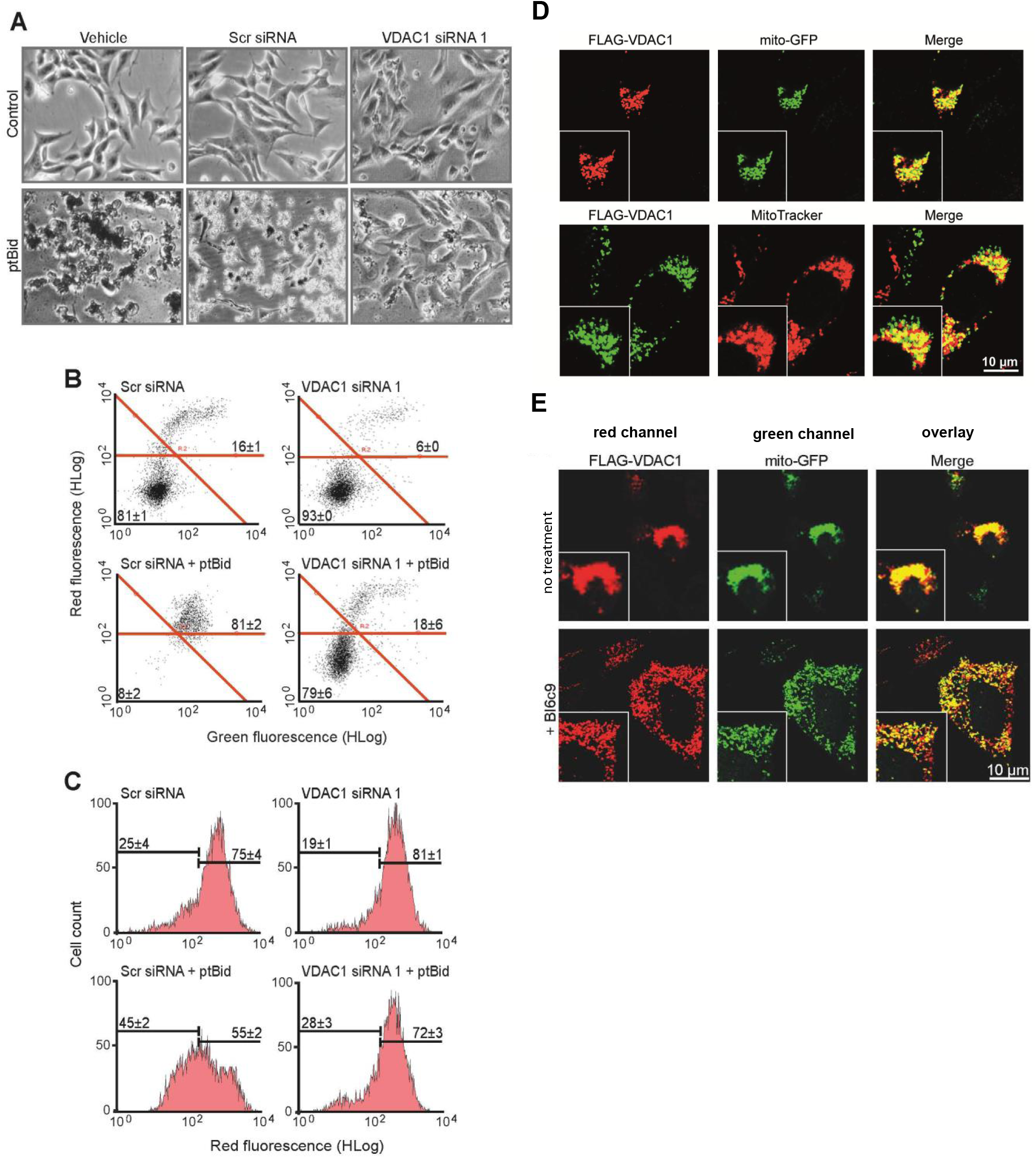
Both VDAC1 and BID are required for neuronal cell death. **(A-C)** HT22 cells were transfected with vehicle, *scr* siRNA, or VDAC1 siRNA 1; 24 h later, the cells were transfected with a tBID-encoding plasmid (ptBID) or mock-transfected (control), and BID toxicity was analyzed 17-18 h later. **(A)** Example images of cells treated as indicated (10x, 0.25-NA objective). **(B)** Annexin V/propidium iodide-FACS analysis of HT22 cells. **(C)** Mitochondrial membrane potential (ΔΨ_m_) was measured using TMRE-FACS. **(D, E)** Localization and effect of VDAC1 at the level of mitochondria. HT22 cells were co-transfected with FLAG-VDAC1 and mito-GFP or transfected with FLAG-VDAC1 and stained with MitoTracker red and then analyzed using confocal fluorescence microscopy. **(D)** Co-localization of VDAC1 (Dylight 648, red upper row and Alexa Fluor 488, green, lower row) with either mito-GFP (upper panel) or MitoTracker (lower panel) indicates that FLAG-tagged VDAC1 protein localizes to the mitochondria (yellow, merge) as also indicated in **E**, upper row. **(E)** VDAC1-induced cytotoxicity is indicated by rounded cells and mitochondria accumulating around the nucleus (**E**, upper row), and this effect is not detected in BI6c9-treated cells, which are preserved from cell death morphology and show mitochondria distributed throughout the cytosol (**E**, lower row). The values in B and C represent the mean ± standard deviation. Images shown in D and E are representative images (n=3-5 experiments with triplicate wells for each condition).

### BID-dependent VDAC1-mediated toxicity

Previous reports showed that overexpression of VDAC1 induces apoptotic cell death [29, 35], and this effect is blocked by inhibiting VDAC channel activity, for example by treating cells with 4,4’-diisothiocyanostilbene-2,2’-disulfonic acid [36], ruthenium red, or hexokinase [35]. However, the mechanism by which VDAC1 overexpression causes cell death has remained poorly understood. Therefore, we investigated whether the putative interaction between BID and VDAC1 is required for VDAC1-mediated cytotoxicity. HT22 cells were co-transfected with FLAG-tagged VDAC1 and analyzed using fluorescence confocal microscopy (Figure 2D, E, S3F). To confirm the mitochondrial localization of the over-expressed FLAG-VDAC1, HT22 cells were either co-transfected with mitochondrial-targeted GFP (mito-GFP) or stained with MitoTracker red specific to visualize the mitochondria (Figure 2D, S3F). Confocal microscopy images indicated a co-localization of both, FLAG-VDAC1 and mito-GFP as well as FLAG-VDAC1 and MitoTracker red, (Figure 2D), suggesting the localization of VDAC1 in the outer mitochondrial membrane (OMM). Furthermore, we investigated whether FLAG-VDAC1 induced overexpression of VDAC1 induced changes in mitochondrial morphology phenotype (Figure S3F). Twenty-four to forty-eight hours after transfection, cells transfected with mito-GFP showed elongated mitochondria (category I) and a spindle-like cell morphology (Figure S3F upper panel). In contrast, cells transfected with FLAG-VDAC1 showed rather fragmented and short mitochondria (mitochondrial morphology category II to category III) and were rounded up (Figure S3F lower panel), indicating that VDAC1 overexpression triggers cytotoxicity through detrimental effects at the level of mitochondria. To further investigate if VDAC1 induced mitochondrial effects are linked to the pro-apoptotic protein BID, cells were co-transfected with FLAG-VDAC1 and mito-GFP after pre-incubation of cells with the BID inhibitor BI6c9 (Figure 2E). As control, and in line with Figure 2D cells co-transfected with FLAG-VDAC1 and mito-GFP, without further treatment, showed a round cell morphology with mitochondria located condensed around the nucleus with a rather dotted morphology (category I) indicating dying cells (Figure 2E, upper panel, no treatment (co-transfection only)). In contrast, pre-incubation of VDAC1-transfected cells with 10 µM of the BID inhibitor BI6c9 prevented mitochondrial morphology changes and thereby cell death (Figure 2E, lower panels). Similar to the findings described before, comparing the expression pattern of FLAG-VDAC1 and mito-GFP confirmed that the recombinant VDAC1 was localized to mitochondrial membranes (Figure 2E, merge). Accordingly, these results demonstrated functional over-expression of recombinant VDAC1 in mitochondria and suggested BID-dependent activation of the mitochondrial VDAC1 leading to cell death.

### Direct physical and functional interactions between BID/tBID and VDAC1 *in vitro*

Our results from the model of glutamate-induced cell death and after overexpression of either tBid or VDAC1 suggest that a direct physical interaction between BID and VDAC1 mediates mitochondrial damage in dying neurons. Therefore, we tested whether BID or its active form tBID bind to VDAC1 by performing *in vitro* pull-down experiments and thermophoresis experiments. We found that purified recombinant VDAC1 (mVDAC1) binds to His_6_-tagged recombinant BID (His-rBID) as well as to His_6_-tagged recombinant tBID (His-rtBID) (Figure3A, S4A and S4B). Further, thermophoresis measurements using labeled mVDAC1 revealed that VDAC1 binds to purified BID and tBID with an apparent dissociation constant (*K*_*D*_) of 1420 ± 3.04 µM and 37.1 ± 1.12 µM, respectively (Figure 3B), values well in line with the stronger apoptotic potency of tBID. This binding was specific for VDAC1, as no binding was measured when BID was incubated with the *Escherichia coli* outer membrane protein G (OmpG) or with empty detergent micelles (Figure 3B).

**Figure 3.**
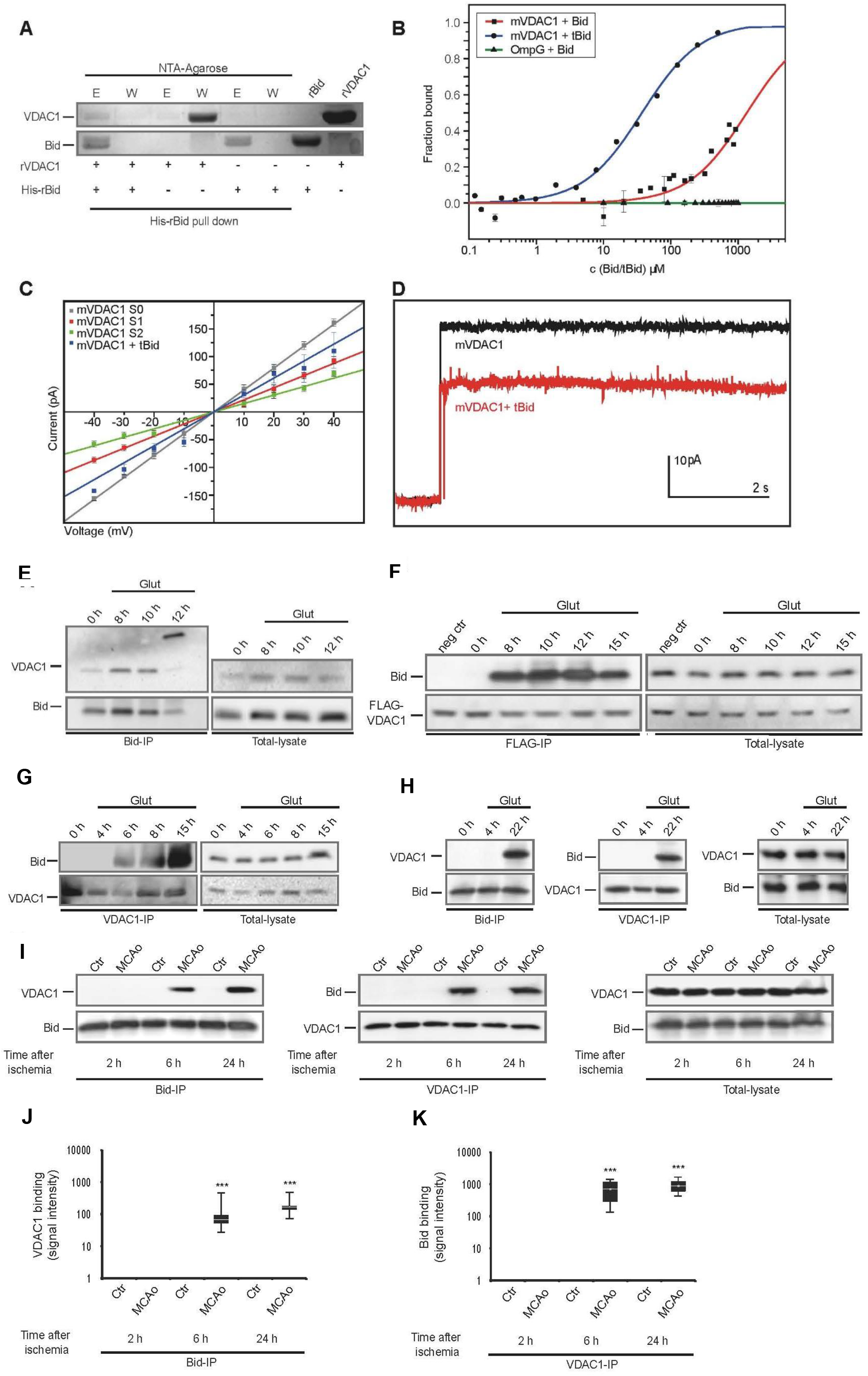
Direct and functional interaction of BID/tBID with VDAC1. **(A)** Pull-down of purified recombinant His_6_-BID incubated with recombinant rVDAC1 as indicated, analyzed by Coomassie-stained 12.5% SDS-PAGE. E: eluted fractions, W: wash fractions. rBID and rVDAC1 were used as protein size controls (last two lanes). **(B)** Thermophoresis measurements were performed under the indicated conditions with fluorescently labeled mVDAC1 or OmpG. The mean ± SD K_D_ values for tBID and BID binding were 37.1±1.12 µM and 1420±3.04 µM, respectively (n=3 independent experiments). **(C)** Current‒voltage plots were used to calculate channel conductance. The S0 (grey), S1 (red), and S2 (green) states had the following calculated conductance: 3.9 ± 0.0, 2.6 ± 0.0, and 1.9 ± 0.1 nS, respectively. In the presence of tBID, channel conductance was 3.0 ± 0.2 nS. The data are presented as the mean ± SE of 10 separate experiments. **(D)** Representative traces of an mVDAC1 channel recorded at +10 mV in the absence (black) or presence (red) of tBID. **(E-G)** Co-immunoprecipitation (left) of VDAC with BID **(E)**, BID with FLAG-VDAC1 **(F)**, or BID with VDAC1 **(G)** in HT22 cells treated with or without 3-5 mM glutamate, followed by western blot analysis of the indicated proteins. Total protein lysates (right) were used as controls. **(H)** Co-immunoprecipitation of BID and VDAC1 (and total protein lysates) from primary cortical neurons following stimulation with 25 µM glutamate. **(I)** Co-immunoprecipitation of BID and VDAC1 from control (ctr) and ischemic brain tissues obtained from mice 2, 6, or 24 h after transient middle cerebral artery occlusion (MCAo). **(J, K)** Summary of the data shown in panel I (n= 6 mice per time point). The upper and lower boxes represent the first and third quartiles, respectively; the horizontal line shows the median; and the upper and lower whiskers indicate the minimum and maximum values, respectively. ***, *p*<0.001 (ANOVA, Kruskal-Wallis *H* test).

Next, we examined the functional role of BID binding on VDAC1 channel activity. We recorded currents through VDAC1 channels in the absence and presence of tBID. VDAC1 channels have both high-conductance (S0) and low-conductance (S1 and S2) states [37–39]. Consistent with previous reports [40], recombinant mVDAC1 channels displayed all three conductance states, with 3.94 ± 0.04 nS, 2.61 ± 0.01 nS, and 1.90 ± 0.06 nS corresponding to the S0, S1, and S2 states, respectively (Figure 3C and S4C). Adding recombinant tBID reduced the channel’s conductance to 3.05 ± 0.24 nS, which represents a 23% decrease relative to the S0 conductance state (Figure 3C and 3D). Furthermore, tBID also induced atypical rapid channel state transitions (Figure S4C). Channels recorded in the presence of tBID did not exhibit S1 or S2 conductance levels, suggesting that binding of tBID to VDAC1 stabilized the channel in a low conductance state.

### BID and VDAC1 interact in dying neurons in vitro and in vivo

We next asked whether we could confirm an interaction between BID and VDAC1 in models of neuronal cell death using both cultured cells and an *in vivo* model of ischemic brain damage. HT22 cells were exposed to x µM glutamate, after which a co-immunoprecipitation (co-IP) assay was performed (Figures 3E-G and S5A-L). We found that BID precipitated together with VDAC1 after cells were exposed to glutamate for 6-12 h. Interestingly, 12 h after onset of glutamate exposure, the VDAC1 antibody detected a band at a size of 70-kDa following pull-down with the BID antibody (Figure 3E), suggesting that BID was bound to VDAC1 dimers that formed at these late time points following the onset of the glutamate treatment. This finding is consistent with previous studies showing that VDAC1 forms dimers and oligomers during apoptosis [41, 42]. Next, we performed the reverse co-IP experiments in HT22 cell lysates, using either endogenous VDAC1 or a FLAG-tagged VDAC1 for the pull-down and BID for the western blot. Both VDAC1 and FLAG-VDAC1 complexes pulled down BID in cell lysates harvested 6 - 15 h after onset of the glutamate exposure (Figures 3F, 3G and S5E-J). As a final test for the *in vitro* association between BID and VDAC1, we performed co-IP experiments in primary cortical neurons following glutamate-induced activation of NMDA receptors to drive excitotoxicity (Figures 3H and S5M-P). Consistent with our results obtained with HT22 cells, the BID-VDAC1 interaction was induced in a time-dependent manner following glutamate stimulation (Figure 5H).

To investigate whether BID and VDAC1 also interact during neuronal cell death *in vivo*, we performed co-IP experiments in brain tissue lysates obtained from mice subjected to focal cerebral ischemia induced by 60 min of middle cerebral artery occlusion (MCAO) followed by reperfusion (Figures 3I-K and S5Q-V). Two, 6, and 24 h after the onset of ischemia, tissue homogenates were prepared from the ischemic cortex and analyzed using co-IP. In control tissue or before neuronal cell death occurs in ischemic cortex, i.e. 2 hours after MCAO, no interaction between BID and VDAC1 was detected. However, 6 and 24 h after the onset of ischemia, time points well known to be associated with post-ischemic neuronal cell death in the cerebral cortex, robust BID-VDAC1 binding was detected (Figure 3I-K). These data strongly suggest that the interaction of BID and VDAC1 occurs not only *in vitro* but also *in vivo* and represents an integral component of neuronal cell death signaling.

## Discussion

Here, we present compelling evidence that BID—and its activated cleavage product, tBID—binds to the mitochondrial protein VDAC1 in neurons, mediating mitochondrial damage and cell death. This interaction was observed at both the physical and functional levels using recombinant proteins, as well as in cellular models of neuronal death and in ischemic brain tissue. Taken together, these data suggest that BID-VDAC1 binding is an essential step in mitochondrial demise and programmed cell death in cultured neurons *in vitro* and in models of neuronal cell death *in vivo*. Our data also shed new light on the mechanisms of BID-dependent intrinsic cell death and the role of VDAC1 in mitochondrial apoptosis; this is particularly relevant, given the controversy associated with the mechanisms that underlie MOMP [19, 43–47]. Here, we show that neither glutamate exposure nor tBID is sufficient to induce mitochondrial dysfunction and cell toxicity in the absence of functional VDAC1; conversely, overexpressing VDAC1 is not sufficient to induce cytotoxicity in the absence of BID. Hence, direct physical interaction of BID and VDAC1, as evidenced by immunoprecipitation, is necessary to trigger mitochondrial damage and cell death in neurons. The current study therefore demonstrates that MOMP-mediated cell death signaling and breakdown of the mitochondrial membrane potential are mediated by the interaction of BID with VDAC1.

So far it was reported that BAX and BAK interact with VDACs. However, since neurons do not express the active, full-length BAK [48, 49], and in non-neuronal cells BAK appears to bind only the VDAC2 isoform [50], BAX and BAK do not seem to be required for tBID-VDAC1 complex formation in neurons. In neurons, the tBID-VDAC1 complex likely does not include BAK. In contrast, BAX is expressed widely throughout the nervous system and is believed to interact with VDAC1; however, the interaction between BAX and VDAC1 for mitochondrial pore formation in living cells has been discussed controversially [17, 51]. Nevertheless, the interaction between tBID and VDAC1 observed here appears to be an essential step in glutamate-induced mitochondrial breakdown and subsequent cell death in neurons, as the removal of either BID or VDAC1 is sufficient to provide neuroprotection. Importantly, cell death pathways may differ between neurons and non-neuronal cells. For example, mouse embryonic fibroblasts that lack VDAC undergo wild-type levels of mitochondrial cytochrome c release and apoptosis [44]; in contrast, reducing the levels of VDAC1 in the brain tissue of mice has neuroprotective effects against lipid peroxidation, ROS production, and mitochondrial dysfunction [52].

With respect to the precise mechanism of this interaction and how it affects downstream signaling cascades, we propose that during cell death stimuli, full-length BID targets VDAC1 to induce a conformational change in the channel. Cleavage of BID and release of the p7 BID cleavage fragment occurs rapidly upon initial binding of cleaved BID to the mitochondrial outer membrane [53] and may strengthen the binding of the remaining tBID fragment to VDAC1. The resulting tBID-VDAC1 complex leads to reduced conductance through the channel and exposes the N-terminal α-helix in VDAC1 to the inner mitochondrial space. As tBID binding reduces the diameter of the channel’s pore, the tBID-VDAC1 complex might reflect a transitional state that further catalyzes the formation of VDAC1 homo-oligomers or hetero-oligomers to sequester a pore formation, enabling the release of death promoting factors into the cytosol and subsequent cell death (Figure 4).

**Figure 4.**
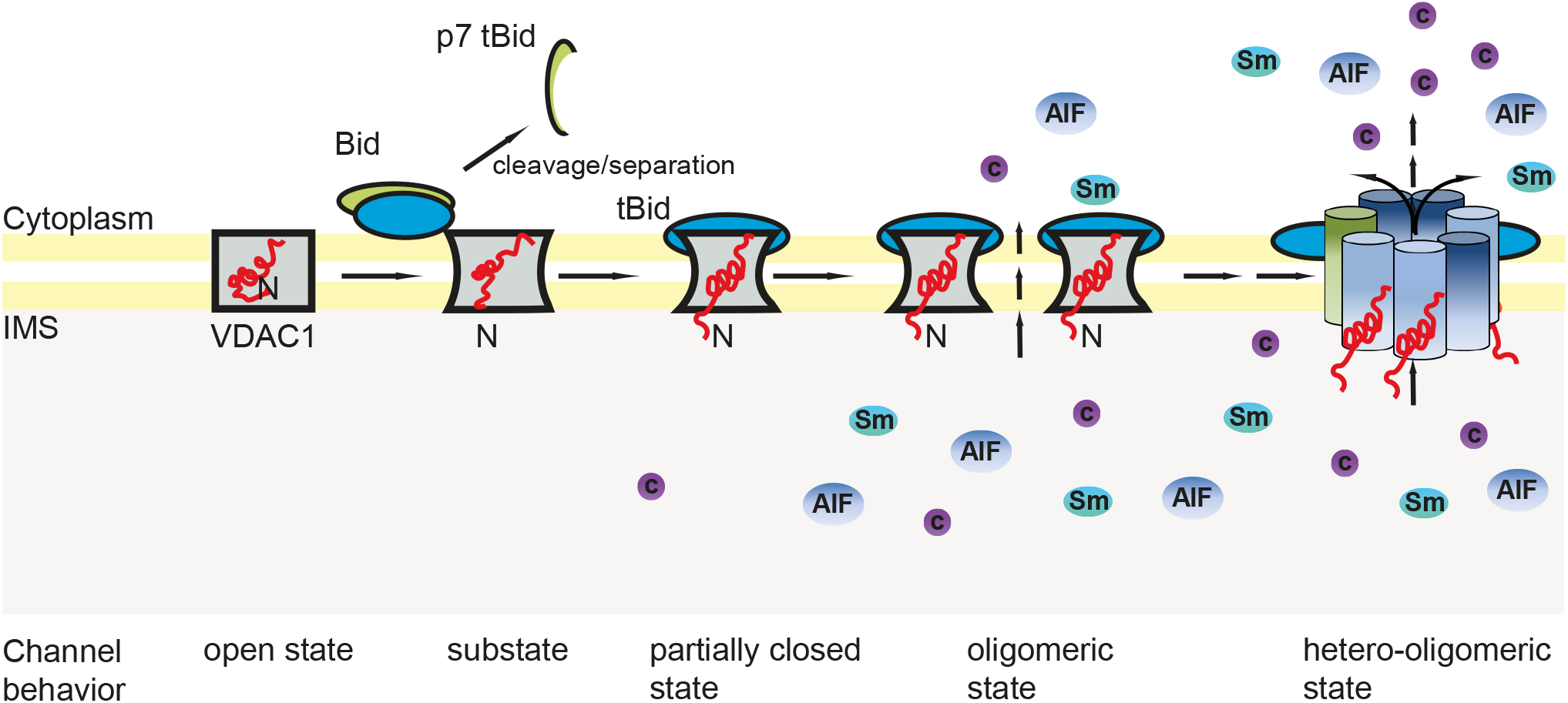
Proposed model for the putative direct interaction between BID and VDAC1 in the mitochondrial membrane. Upon receiving an apoptotic stimulus, BID and VDAC1 form a complex. The association of full-length BID with VDAC1 induces a conformational change in VDAC1 and facilitates the cleavage of BID and release of the p7 tBID fragment. The remaining p15 tBID fragment (indicated as “tBID”) binds tightly to VDAC1 and is oriented parallel to the membrane surface [54]. The tBID-VDAC1 complex might reflect a transitional state with a reduced overall current that enables the exposure of the VDAC1 α-helix to the inner mitochondrial space (IMS). This partially closed state of VDAC1 together with the exposed α-helix, catalyzes the further formation of VDAC1 oligomers or hetero-oligomers to form a pore diameter that enables the release of death promoting factors, such as apoptosis inducing factor (AIF), cytochrome c (c) and Smac/DIABLO (Sm), into the cytosol and the subsequent cell death.

In conclusion, we provide evidence that the protein channel VDAC1 acts as a mitochondrial receptor for BID during neuronal cell death. This interaction may support MOMP, which was previously attributed solely to either the activity of apoptogenic BH3-only proteins or VDACs, respectively. Targeting the interaction between full length BID or tBID and VDAC1 will likely have high therapeutic potential in treating neurological conditions in which mitochondrial cell death pathways play a prominent role.

## Author contributions

S.O. designed and performed the experiments, analyzed the experiments, and wrote the manuscript. L.M., S.O. and performed the *in vivo* experiments; L.M. helped writing the manuscript; B.M., G.P., produced the recombinant VDAC1, performed the thermophoresis analysis, and performed the lipid bilayer experiments together with P.R.; C.K. and. generated the FLAG-tagged VDAC1 plasmid; A.M.D. performed the Seahorse Bioscience measurements. L.O.W., M.B. supervised experiments and edited the manuscript. N.P. supervised experiments and edited figures and manuscript, C.C. contributed to study concept and design, supervised the study, wrote and edited the manuscript. All authors have read and approved the manuscript.

## Acknowledgments

We thank Dr. Jeff Abramson (UCLA) for kindly providing the mVDAC1 plasmid and Prof. Jean-Claude Martinou (University of Geneva, Switzerland) for kindly providing recombinant caspase 8. We are grateful to Uta Mamrak, Katharina Elsässer, and Eileen Daube for providing technical support. We also thank the late Prof. Walter Neupert for commenting on the manuscript. We also thank Dr. Curtis Barrett (English Editing Solutions) for editing the manuscript and for providing comments and suggestions. Part of this work was supported by a grant of the Deutsche Forschungsgemeinschaft and the Von-Behring-Röntgen Foundation to CC (DFG FOR2107 CU 43/9-1; CU 43/12-1, BR62-0023).

## References

1. Galluzzi L, Vitale I, Abrams JM, Alnemri ES, Baehrecke EH, Blagosklonny MV, et al. Molecular definitions of cell death subroutines: recommendations of the Nomenclature Committee on Cell Death 2012. Cell Death Differ. 2012;19:107–20. doi:10.1038/cdd.2011.96.

2. Kroemer G, Galluzzi L, Brenner C. Mitochondrial Membrane Permeabilization in Cell Death. Physiological Reviews. 2007;87:99–163. doi:10.1152/physrev.00013.2006.

3. Zhu C, Qiu L, Wang X, Hallin U, Candé C, Kroemer G, et al. Involvement of apoptosis-inducing factor in neuronal death after hypoxia-ischemia in the neonatal rat brain. Journal of Neurochemistry. 2003;86:306–17. doi:10.1046/j.1471-4159.2003.01832.x.

4. Chan DC. Mitochondria: Dynamic Organelles in Disease, Aging, and Development. Cell. 2006;125:1241–52. doi:10.1016/j.cell.2006.06.010.

5. Culmsee C, Landshamer S. Molecular Insights into Mechanisms of the Cell Death Program:Role in the Progression of Neurodegenerative Disorders. CAR. 2006;3:269–83. doi:10.2174/156720506778249461.

6. Darwish R. Regulatory mechanisms of apoptosis in regularly dividing cells. CHC. 2010:59. doi:10.2147/CHC.S10036.

7. Chipuk JE, Bouchier-Hayes L, Green DR. Mitochondrial outer membrane permeabilization during apoptosis: the innocent bystander scenario. Cell Death Differ. 2006;13:1396–402. doi:10.1038/sj.cdd.4401963.

8. Desagher S, Martinou J-C. Mitochondria as the central control point of apoptosis. Trends in Cell Biology. 2000;10:369–77. doi:10.1016/S0962-8924(00)01803-1.

9. Green DR. The Pathophysiology of Mitochondrial Cell Death. Science. 2004;305:626–9. doi:10.1126/science.1099320.

10. Shoshan-Barmatz V, Ben-Hail D, Admoni L, Krelin Y, Tripathi SS. The mitochondrial voltage-dependent anion channel 1 in tumor cells. Biochimica et Biophysica Acta (BBA) - Biomembranes. 2015;1848:2547–75. doi:10.1016/j.bbamem.2014.10.040.

11. Dadsena S, Bockelmann S, Mina JGM, Hassan DG, Korneev S, Razzera G, et al. Ceramides bind VDAC2 to trigger mitochondrial apoptosis. Nat Commun 2019. doi:10.1038/s41467-019-09654-4.

12. Shoshan-Barmatz V, Golan M. Mitochondrial VDAC1: Function in Cell Life and Death and a Target for Cancer Therapy. CMC. 2012;19:714–35. doi:10.2174/092986712798992110.

13. Tsujimoto Y, Shimizu S. The voltage-dependent anion channel: an essential player in apoptosis. Biochimie. 2002;84:187–93. doi:10.1016/S0300-9084(02)01370-6.

14. Colombini M. VDAC: The channel at the interface between mitochondria and the cytosol. Mol Cell Biochem. 2004;256:107–15. doi:10.1023/B:MCBI.0000009862.17396.8d.

15. Lemasters JJ, Holmuhamedov E. Voltage-dependent anion channel (VDAC) as mitochondrial governator—Thinking outside the box. Biochimica et Biophysica Acta (BBA) - Molecular Basis of Disease. 2006;1762:181–90. doi:10.1016/j.bbadis.2005.10.006.

16. Nagakannan P, Islam MI, Karimi-Abdolrezaee S, Eftekharpour E. Inhibition of VDAC1 Protects Against Glutamate-Induced Oxytosis and Mitochondrial Fragmentation in Hippocampal HT22 Cells. Cell Mol Neurobiol. 2019;39:73–85. doi:10.1007/s10571-018-0634-1.

17. Rostovtseva TK, Antonsson B, Suzuki M, Youle RJ, Colombini M, Bezrukov SM. Bid, but Not Bax, Regulates VDAC Channels. Journal of Biological Chemistry. 2004;279:13575–83. doi:10.1074/jbc.M310593200.

18. Banerjee J, Ghosh S. Bax increases the pore size of rat brain mitochondrial voltage-dependent anion channel in the presence of tBid. Biochemical and Biophysical Research Communications. 2004;323:310–4. doi:10.1016/j.bbrc.2004.08.094.

19. Shimizu S, Shinohara Y, Tsujimoto Y. Bax and Bcl-xL independently regulate apoptotic changes of yeast mitochondria that require VDAC but not adenine nucleotide translocator. Oncogene. 2000;19:4309–18. doi:10.1038/sj.onc.1203788.

20. Tsujimoto Y, Shimizu S. VDAC regulation by the Bcl-2 family of proteins. Cell Death Differ. 2000;7:1174–81. doi:10.1038/sj.cdd.4400780.

21. Shoshan-Barmatz V, Keinan N, Abu-Hamad S, Tyomkin D, Aram L. Apoptosis is regulated by the VDAC1 N-terminal region and by VDAC oligomerization: release of cytochrome c, AIF and Smac/Diablo. Biochimica et Biophysica Acta (BBA) - Bioenergetics. 2010;1797:1281–91. doi:10.1016/j.bbabio.2010.03.003.

22. Plesnila N, Zinkel S, Le DA, Amin-Hanjani S, Wu Y, Qiu J, et al. BID mediates neuronal cell death after oxygen/ glucose deprivation and focal cerebral ischemia. Proceedings of the National Academy of Sciences. 2001;98:15318–23. doi:10.1073/pnas.261323298.

23. Landshamer S, Hoehn M, Barth N, Duvezin-Caubet S, Schwake G, Tobaben S, et al. Bid-induced release of AIF from mitochondria causes immediate neuronal cell death. Cell Death Differ. 2008;15:1553–63. doi:10.1038/cdd.2008.78.

24. Grohm J, Kim S-W, Mamrak U, Tobaben S, Cassidy-Stone A, Nunnari J, et al. Inhibition of Drp1 provides neuroprotection in vitro and in vivo. Cell Death Differ. 2012;19:1446–58. doi:10.1038/cdd.2012.18.

25. Neitemeier S, Jelinek A, Laino V, Hoffmann L, Eisenbach I, Eying R, et al. BID links ferroptosis to mitochondrial cell death pathways. Redox Biology. 2017;12:558–70. doi:10.1016/j.redox.2017.03.007.

26. Diemert S, Dolga AM, Tobaben S, Grohm J, Pfeifer S, Oexler E, Culmsee C. Impedance measurement for real time detection of neuronal cell death. Journal of Neuroscience Methods. 2012;203:69–77. doi:10.1016/j.jneumeth.2011.09.012.

27. Jelinek A, Heyder L, Daude M, Plessner M, Krippner S, Grosse R, et al. Mitochondrial rescue prevents glutathione peroxidase-dependent ferroptosis. Free Radic Biol Med. 2018;117:45–57. doi:10.1016/j.freeradbiomed.2018.01.019.

28. Tobaben S, Grohm J, Seiler A, Conrad M, Plesnila N, Culmsee C. Bid-mediated mitochondrial damage is a key mechanism in glutamate-induced oxidative stress and AIF-dependent cell death in immortalized HT-22 hippocampal neurons. Cell Death Differ. 2011;18:282–92. doi:10.1038/cdd.2010.92.

29. Abu-Hamad S, Sivan S, Shoshan-Barmatz V. The expression level of the voltage-dependent anion channel controls life and death of the cell. Proceedings of the National Academy of Sciences. 2006;103:5787–92. doi:10.1073/pnas.0600103103.

30. Grohm J, Plesnila N, Culmsee C. Bid mediates fission, membrane permeabilization and perinuclear accumulation of mitochondria as a prerequisite for oxidative neuronal cell death. Brain, Behavior, and Immunity. 2010;24:831–8. doi:10.1016/j.bbi.2009.11.015.

31. Öxler E-M, Dolga A, Culmsee C. AIF depletion provides neuroprotection through a preconditioning effect. Apoptosis. 2012;17:1027–38. doi:10.1007/s10495-012-0748-8.

32. Culmsee C, Plesnila N. Targeting Bid to prevent programmed cell death in neurons. Biochemical Society Transactions. 2006;34:1334–40. doi:10.1042/BST0341334.

33. Barho MT, Oppermann S, Schrader FC, Degenhardt I, Elsässer K, Wegscheid-Gerlach C, et al. N -Acyl Derivatives of 4-Phenoxyaniline as Neuroprotective Agents. ChemMedChem. 2014;9:2260–73. doi:10.1002/cmdc.201402195.

34. Oppermann S, Schrader FC, Elsässer K, Dolga AM, Kraus AL, Doti N, et al. Novel N -Phenyl– Substituted Thiazolidinediones Protect Neural Cells against Glutamate-and tBid-Induced Toxicity. J Pharmacol Exp Ther. 2014;350:273–89. doi:10.1124/jpet.114.213777.

35. Zaid H, Abu-Hamad S, Israelson A, Nathan I, Shoshan-Barmatz V. The voltage-dependent anion channel-1 modulates apoptotic cell death. Cell Death Differ. 2005;12:751–60. doi:10.1038/sj.cdd.4401599.

36. Godbole A, Varghese J, Sarin A, Mathew MK. VDAC is a conserved element of death pathways in plant and animal systems. Biochimica et Biophysica Acta (BBA) - Molecular Cell Research. 2003;1642:87–96. doi:10.1016/S0167-4889(03)00102-2.

37. Abu-Hamad S, Arbel N, Calo D, Arzoine L, Israelson A, Keinan N, et al. The VDAC1 N-terminus is essential both for apoptosis and the protective effect of anti-apoptotic proteins. Journal of Cell Science. 2009;122:1906–16. doi:10.1242/jcs.040188.

38. Hiller S, Garces RG, Malia TJ, Orekhov VY, Colombini M, Wagner G. Solution Structure of the Integral Human Membrane Protein VDAC-1 in Detergent Micelles. Science. 2008;321:1206–10. doi:10.1126/science.1161302.

39. Ujwal R, Cascio D, Colletier J-P, Faham S, Zhang J, Toro L, et al. The crystal structure of mouse VDAC1 at 2.3 A resolution reveals mechanistic insights into metabolite gating. Proc Natl Acad Sci U S A. 2008;105:17742–7. doi:10.1073/pnas.0809634105.

40. Mertins B, Psakis G, Grosse W, Back KC, Salisowski A, Reiss P, et al. Flexibility of the N-Terminal mVDAC1 Segment Controls the Channel’s Gating Behavior. PLoS ONE. 2012;7:e47938. doi:10.1371/journal.pone.0047938.

41. Geula S, Naveed H, Liang J, Shoshan-Barmatz V. Structure-based Analysis of VDAC1 Protein. Journal of Biological Chemistry. 2012;287:2179–90. doi:10.1074/jbc.M111.268920.

42. Mader A, Abu-Hamad S, Arbel N, Gutierr-Asuilar M, Shoshan-Barmatz V. Dominant-negative VDAC1 mutants reveal oligomeric VDAC1 to be the active unit in mitochondria-mediated apoptosis. Biochemical Journal. 2010;429:147–55. doi:10.1042/BJ20091338.

43. Arbel N, Ben-Hail D, Shoshan-Barmatz V. Mediation of the Antiapoptotic Activity of Bcl-xL Protein upon Interaction with VDAC1 Protein. Journal of Biological Chemistry. 2012;287:23152–61. doi:10.1074/jbc.M112.345918.

44. Baines CP, Kaiser RA, Sheiko T, Craigen WJ, Molkentin JD. Voltage-dependent anion channels are dispensable for mitochondrial-dependent cell death. Nat Cell Biol. 2007;9:550–5. doi:10.1038/ncb1575.

45. Rostovtseva TK, Bezrukov SM. VDAC regulation: role of cytosolic proteins and mitochondrial lipids. J Bioenerg Biomembr. 2008;40:163–70. doi:10.1007/s10863-008-9145-y.

46. Shimizu S, Narita M, Tsujimoto Y. Bcl-2 family proteins regulate the release of apoptogenic cytochrome c by the mitochondrial channel VDAC. Nature. 1999;399:483–7. doi:10.1038/20959.

47. Shimizu S, Tsujimoto Y. Proapoptotic BH3-only Bcl-2 family members induce cytochrome c release, but not mitochondrial membrane potential loss, and do not directly modulate voltage-dependent anion channel activity. Proceedings of the National Academy of Sciences. 2000;97:577–82. doi:10.1073/pnas.97.2.577.

48. Jakobson M, Lintulahti A, Arumäe U. mRNA for N-Bak, a neuron-specific BH3-only splice isoform of Bak, escapes nonsense-mediated decay and is translationally repressed in the neurons. Cell Death Dis. 2012;3:e269. doi:10.1038/cddis.2012.4.

49. Jakobson M, Llano O, Palgi J, Arumäe U. Multiple mechanisms repress N-Bak mRNA translation in the healthy and apoptotic neurons. Cell Death Dis. 2013;4:e777. doi:10.1038/cddis.2013.297.

50. Chandra D, Choy G, Daniel PT, Tang DG. Bax-dependent Regulation of Bak by Voltage-dependent Anion Channel 2. Journal of Biological Chemistry. 2005;280:19051–61. doi:10.1074/jbc.M501391200.

51. Keinan N, Tyomkin D, Shoshan-Barmatz V. Oligomerization of the Mitochondrial Protein Voltage-Dependent Anion Channel Is Coupled to the Induction of Apoptosis. Mol Cell Biol. 2010;30:5698–709. doi:10.1128/MCB.00165-10.

52. Manczak M, Sheiko T, Craigen WJ, Reddy PH. Reduced VDAC1 Protects Against Alzheimer’s Disease, Mitochondria, and Synaptic Deficiencies. JAD. 2013;37:679–90. doi:10.3233/JAD-130761.

53. Shamas-Din A, Bindner S, Zhu W, Zaltsman Y, Campbell C, Gross A, et al. tBid Undergoes Multiple Conformational Changes at the Membrane Required for Bax Activation. Journal of Biological Chemistry. 2013;288:22111–27. doi:10.1074/jbc.M113.482109.

54. Wang Y, Tjandra N. Structural Insights of tBid, the Caspase-8-activated Bid, and Its BH3 Domain. Journal of Biological Chemistry. 2013;288:35840–51. doi:10.1074/jbc.M113.503680.

