## Supplemental material for "Interaction between BID and VDAC1 is required for mitochondrial demise and cell death in neurons"

**Supplemental Figure legends**

**Figure S1, related to Figure 1: Silencing of VDAC1 attenuates glutamate-induced toxicity in HT-22 cells. (A)**, **(B)** VDAC1 knockdown by two different VDAC1 siRNA sequences (VDAC1 siRNA 1, VDAC1 siRNA 2) verified by RT-PCR analysis of VDAC1 mRNA **(A)** and western blot analysis of VDAC1 protein **(B)** 48 h after application of 20 nM VDAC1 siRNA 1 or siRNA 2. Nonfunctional scrambled (*scr*) siRNA or Lipofectamine RNAimax (vehicle) was applied in control experiments. RT-PCR with primers specific for glyceraldehyde-3-phosphatedehydrogenase (GAPDH) and anti-β-Actin antibodies were used as controls for respective analysis. **(C)** RT-PCR analysis of VDAC2 mRNA levels ruled out unspecific knockdown by VDAC1 siRNA 1 and siRNA 2 in concentrations of 20 nM to 80 nM. **(D)** Photomicrographs (10 x 0.25 NA objective) of HT-22 cells reveal protection against glutamate-induced (3 mM, 16 h) cytotoxicity by VDAC1 siRNA 2. **(E)** FACS analysis of annexin-V/propidium iodide (AV/PI) stained HT-22 cells 17.5 h after glutamate exposure (5 mM). Numbers are mean percentages ± s.d. for three cell groups treated as indicated in the corresponding quadrants. Lower left corner indicates AV-/PI-healthy cells; upper right corner depicts AV+/PI+ dead cells. **(F)** Quantification of AV/PI-FACS analysis **(E)** confirmed significant inhibition of cell death by both VDAC1 siRNAs. **(G)** MTT assay confirmed the protective effect of VDAC1 siRNA 2 (20 nM) against glutamate toxicity (3 mM, 16 h). (***p < 0.001 compared to glutamate treated vehicle and scr siRNA, ANOVA, Scheffe´test **(F)**, **(G)**). **(H)** Real-time detection of cellular impedance by xCELLigence System (Roche, Penzberg, Germany). VDAC1 siRNA 2 transfected cells (dark green circle) are similarly protected against glutamate- induced cell death (Glut, 4 mM) as VDAC1 siRNA 1-treated cells (Figure 1). For statistical analysis experiments were independently repeated three to five times with n=3 **(A-G)** or n=8 **(D)** per treatment condition and data are provided as mean±s.d. Glut, Glutamate.

**Figure S2, related to Figure 1: VDAC1 siRNA 2 prevents glutamate-induced mitochondrial integrity and function in HT-22 cells. (A)** BODIPY-FACS analysis of lipid peroxidation 18 h after glutamate treatment (4 mM) of HT-22 cells. VDAC1 siRNA 2 prevents glutamate-induced lipid peroxidation. Values in the corresponding quadrants are averages (%)±s.d for three indicated treatment groups. **(B)** Quantification of **(A)** showed significant prevention of lipid peroxidation by both VDAC1siRNAs. **(C)** ATP-luminescence measurements of HT-22 cells, transfected as indicated, in the presence or absence of glutamate (5 mM, 20 h). VDAC1 siRNA 2 prevents glutamate-inducedATP depletion. **(D)** Seahorse Bioscience measurements of oxygen consumption rate (OCR). VDAC1 siRNA 2 preserves mitochondrial maximum respiration and respiratory capacity after glutamate exposure of HT-22 cells. **(G)** Confocal fluorescence photomicrographs reveal protection against glutamate-induced mitochondrial fragmentation in VDAC1 siRNA 2 (20 nM) transfected HT-22 cells. Mitochondria were visualized by Mitotracker red staining 17 h after glutamate treatment (5 mM). **(E)** TMRE-FACS analysis of mitochondrial membrane potential of HT-22 cells. VDAC1siRNA 2 prevents the glutamate-induced (17 h, 4 mM) drop in red fluorescence (loss of$\Delta\psi_m$). Numbers are mean percentages±s.d. of TMRE fluorescence for n=3 indicated groups. **(F)** Quantification of TMRE fluorescence **(E)** confirms the preservation of mitochondrial membrane potential by VDAC1 gene silencing using two different VDAC1 siRNA sequences (VDAC1 siRNA1 and VDAC1 siRNA2). **(H)** Quantification of mitochondrial morphology: Category 1: elongated, category II:intermediate, category III: fragmented mitochondria. Data represent the mean±s.d. of three independent experiments (###p < 0.001 compared to category I glutamate treated control and *scr*siRNA; ***p < 0.001 compared to category III glutamate treated control and scr siRNA, ANOVA, Scheffe´ test). **(A-H)** All experiment were independently repeated at least three to five times and data are provided as mean±s.d. (###p < 0.001 compared to vehicle and *scr* siRNA controls, ***p <0.001 compared to glutamate treated vehicle and scr siRNA, n=6-8, ANOVA, Scheffé test). Glut, Glutamate.

**Figure S3, related to Figure 2: tBid-induced toxicity and loss of $\Delta\psi_m$ are prevented by VDAC1 gene silencing.** HT-22 cells transfected with either VDAC1 siRNA 1 or VDAC1 siRNA 2 were posttransfected (24 h) with a tBid encoding plasmid (ptBid). Cells treated only with attractene (vehicle) or *scr* siRNA transfected cells were used as controls. **(A)** VDAC1 siRNA 2 preserves cell morphology 17 h after tBid over-expression (ptBid) (photomicroscopy; 10 x 0.25 NA objective). **(B)** FACS recordings of annexin-V/propidium iodide stained HT-22 cells showed resistance of VDAC1 siRNA 2 transfected cells to tBid-induced cell death. Numbers are mean values (%)±s.d. for three indicated treatment groups in the corresponding quadrants. **(C)** Quantification of AV+/PI+ cells confirmed that VDAC1-depleted cells (VDAC1 siRNA 1and VDAC1 siRNA 2) are protected against tBid-induced cell death. **(D)** TMRE-FACS measurements of mitochondrial membrane potential 18 h after tBid overexpression. Numbers are mean percentages±s.d. of n=3 per treatment group, indicating loss of $\Delta\psi_m$ (left side) or intact $\Delta\psi_m$ (right side). **(E)** Quantification of TMRE fluorescence revealed significant loss of $\Delta\psi_m$ in HT-22 cells over-expressing tBid which was prevented by both VDAC1 siRNAs. **(B-E)** ***$p < 0.001$ compared to pIRES-tBid transfected vehicle and *scr* siRNA, n=3,ANOVA, Scheffé test. All experiments were independently repeated at least three times. (**F)** Confocal microscopy of HT22 cells transfected with either mito-GFP (upper row) or FLAG-VDAC1 (lower row), indicating elongated mitochondria (category I) and spindle-like cells upper row and a VDAC1-induced fragmentation of mitochondria surrounding the nucles in FLAG-VDAC1 tranfected cells (lower row).

**Figure S4, related to Figure 3: Direct and functional interaction of Bid/tBid andVDAC1.** **(A)** Direct interaction between purified rBid and rVDAC1. Original and un-cut coomassie stained 12.5% SDS-PAGE is shown. E, elution-fraction; W, wash-fraction. **(B)** The direct interaction between purified rBid and rVDAC1 was confirmed by Western blot analysis. For generation of tBid, recombinant full-length Bid was incubated with recombinant caspase 8 for 2 h at RT. Elution-fractions (E) and wash-fractions (W) of the pull down experiments were analyzed by SDS-PAGE followed by western blot. Anti-Bid antibody was used to detect full-length Bid and tBid (upper panel). Binding of VDAC1 to full-length Bid and tBid was indicated by VDAC1 bands in His-rBid and His-rtBid elution fractions. For comparison, deletion of the N-terminal helix of VDAC1 (rVDAC1 Δ11) did not affect co-pull down with Bid. **(C)** BLM-measurements of reconstituted VDAC1. A: representative trace of the mVDAC1 gating activity at +40 mV. The observed S0, S1 and S2 states in the + 40 mV trace are indicated by blue, red and green lines, respectively. B: representative trace of mVDAC1 after addition of tBid at +40 mV. The channel shows minor yet atypical fast channel switching events.

**Figure S5, related to Figure 3: Direct interaction of Bid with VDAC1 in cultured neurons and in ischemic brain tissue of mice.** **(A-H)** Direct interaction of endogenous Bid with endogenous VDAC1 in HT-22 cells exposed to glutamate (3-5 mM) for 4 h to 15 h. **(A)**, **(B)** Bid immune complexes detected with anti-Bid antibody **(A)** and anti-VDAC1 antibody **(B)** and western blots of the corresponding total protein lysates showing Bid **(C)** and VDAC1 **(D)**. The detected lines in the upper half of the anti-Bid pull down western blot **(A)** indicate heavy and light chain of the anti-Bid antibody (rabbit) used for immunoprecipitation. **(E-H)** VDAC1 immune complexes **(E,F)** and corresponding total protein lysates **(G,H)** western-blotted with anti-Bid antibody **(E,G)** and anti-VDAC1 antibody **(F,H)**. Six hours after glutamate exposure a pronounced binding of Bid to VDAC1 was detected, this is increased up to 15 h glutamate treatment **(E)**.**(I-L)** Immunoprecipitation of Flag-VDAC1 over-expressed in HT-22 cells. Flag-VDAC1immune complexes reveal interaction with Bid 8 h to 15 h after treatment with 5 mM glutamate shown by western blot with anti-Bid antibody **(I)**. FLAG-VDAC1 was detected with the anti-Flag antibody **(F)**. Western blots for corresponding total protein lysates detecting Bid **(K)** and Flag-VDAC1 **(L)**. **(M-P)** Immunoprecipitation of endogenous Bid and endogenous VDAC1 from primary cortical neurons 0 h to 22 h after glutamate-induced excitotoxicity. Bid- and VDAC1-immune complexes were western-blotted with anti-Bid antibody **(M)** and anti-VDAC1 antibody **(N)**. Corresponding total protein lysates from PCN were western-blotted with anti-Bid antibody **(O)** and anti-VDAC1 antibody **(P)**. Glut, Glutamate. **(Q-V)** Immunoprecipitation of endogenous Bid and endogenous VDAC1 in brain tissue of mice 2 h, 6 h and 24 h after transient middle cerebral artery occlusion (MCAo). Contralateral brain tissue was used as corresponding control (ctr) to ischemic brain tissue (ipsilateral, MCAo) at the indicated time-points. **(Q)**, **(R)** Bid immune complexes were western-blotted with anti-VDAC1 antibody **(Q)** and anti-Bid antibody **(R)**. **(S)**, **(T)** VDAC1 immune complexes, western-blotted with anti-Bid antibody **(S)** and anti-VDAC1 antibody **(T)** confirm the direct interaction between Bid and VDAC1 in brain tissue6 h and 24 h after ischemia. Corresponding total protein lysates confirm the expression ofBid **(U)** and VDAC1 **(V)** in contralateral control (ctr) and ischemic brain tissue (MCAo).

**Figure S1**

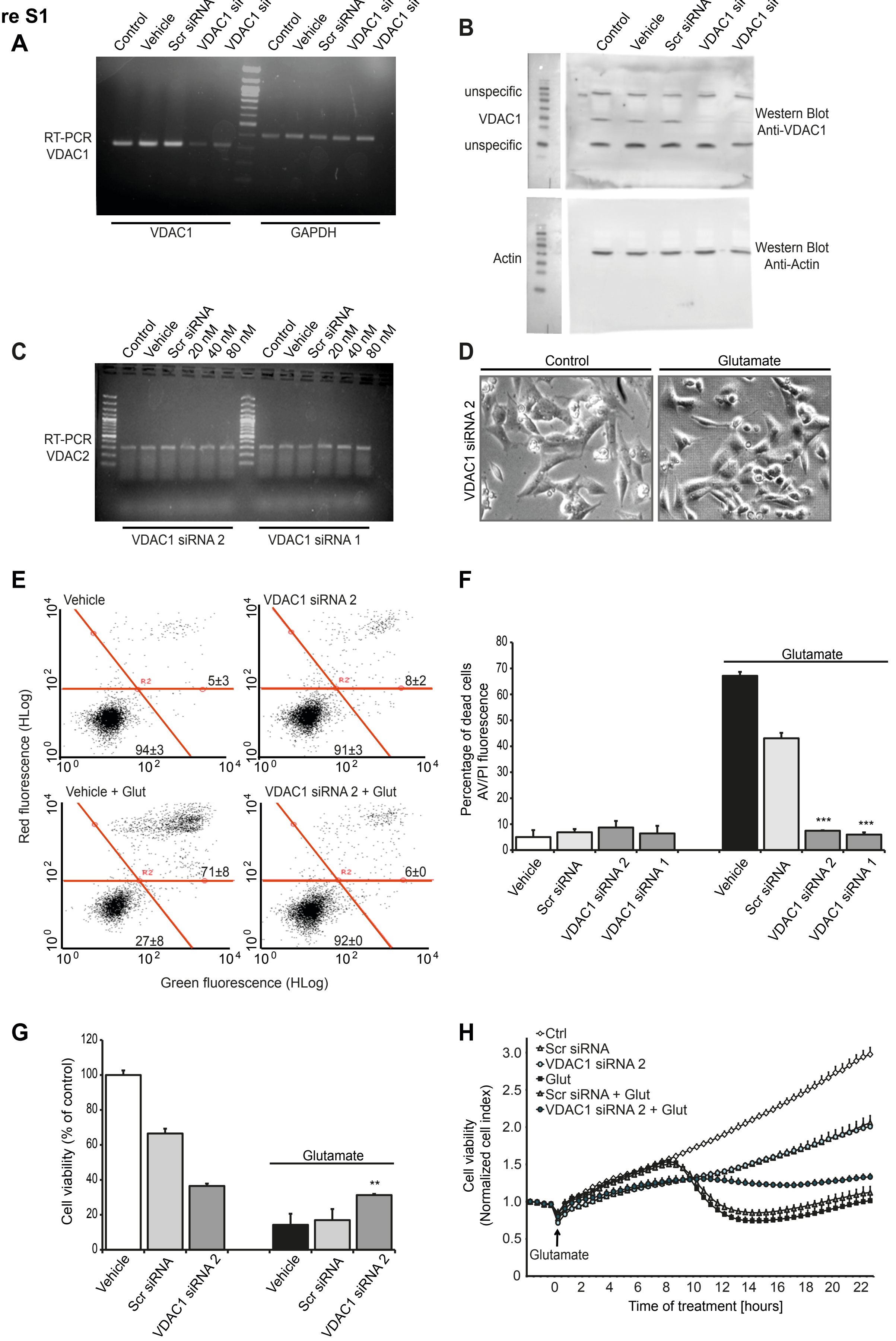

A
Control
Vehicle
Scr siRNA
RT-PCR
VDAC1
VDAC1
GAPDH
B
Control
Vehicle
Scr siRNA
unspecific
VDAC1
unspecific
Western Blot
Anti-VDAC1
Actin
Western Blot
Anti-Actin
C
Control
Vehicle
Scr siRNA
20 nM
40 nM
80 nM
RT-PCR
VDAC2
VDAC1 siRNA 2
VDAC1 siRNA 1
D
Control
Glutamate
VDAC1 siRNA 2
E
Vehicle
VDAC1 siRNA 2
Vehicle + Glut
VDAC1 siRNA 2 + Glut
5±3
94±3
8±2
91±3
71±8
27±8
6±0
92±0
Red fluorescence (HLog)
Green fluorescence (HLog)
F
Percentage of dead cells
AV/PI fluorescence
Glutamate
Vehicle
Scr siRNA
VDAC1 siRNA 2
VDAC1 siRNA 1
***
G
Cell viability (% of control)
Glutamate
Vehicle
Scr siRNA
VDAC1 siRNA 2
**
H
Ctrl
Scr siRNA
VDAC1 siRNA 2
Glut
Scr siRNA + Glut
VDAC1 siRNA 2 + Glut
Cell viability
(Normalized cell index)
Glutamate
Time of treatment [hours]

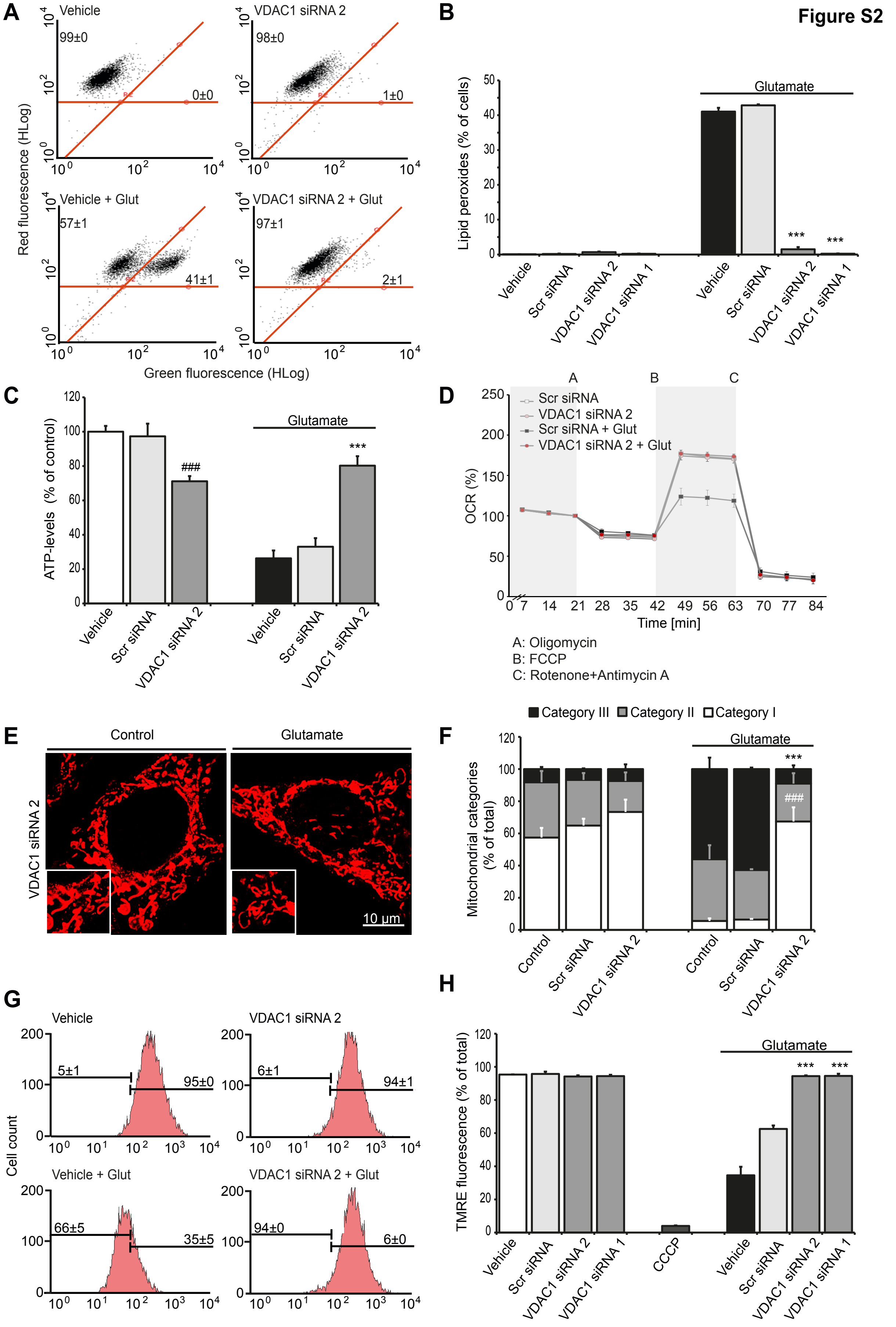

Figure S2
A
Vehicle
99±0
0±0
VDAC1 siRNA 2
98±0
1±0
Vehicle + Glut
57±1
41±1
VDAC1 siRNA 2 + Glut
97±1
2±1
Red fluorescence (HLog)
Green fluorescence (HLog)
B
Lipid peroxides (% of cells)
Glutamate
***
***
Vehicle
Scr siRNA
VDAC1 siRNA 2
VDAC1 siRNA 1
C
ATP-levels (% of control)
Glutamate
###
***
Vehicle
Scr siRNA
VDAC1 siRNA 2
D
A
B
C
Scr siRNA
VDAC1 siRNA 2
Scr siRNA + Glut
VDAC1 siRNA 2 + Glut
OCR (%)
Time [min]
A: Oligomycin
B: FCCP
C: Rotenone+Antimycin A
E
Control
Glutamate
VDAC1 siRNA 2
10 µm
F
Category III
Category II
Category I
Glutamate
***
###
Mitochondrial categories (% of total)
Control
Scr siRNA
VDAC1 siRNA 2
G
Vehicle
5±1
95±0
VDAC1 siRNA 2
6±1
94±1
Vehicle + Glut
66±5
35±5
VDAC1 siRNA 2 + Glut
94±0
6±0
Cell count
H
TMRE fluorescence (% of total)
Glutamate
***
***
Vehicle
Scr siRNA
VDAC1 siRNA 2
VDAC1 siRNA 1
CCCP

Figure S3

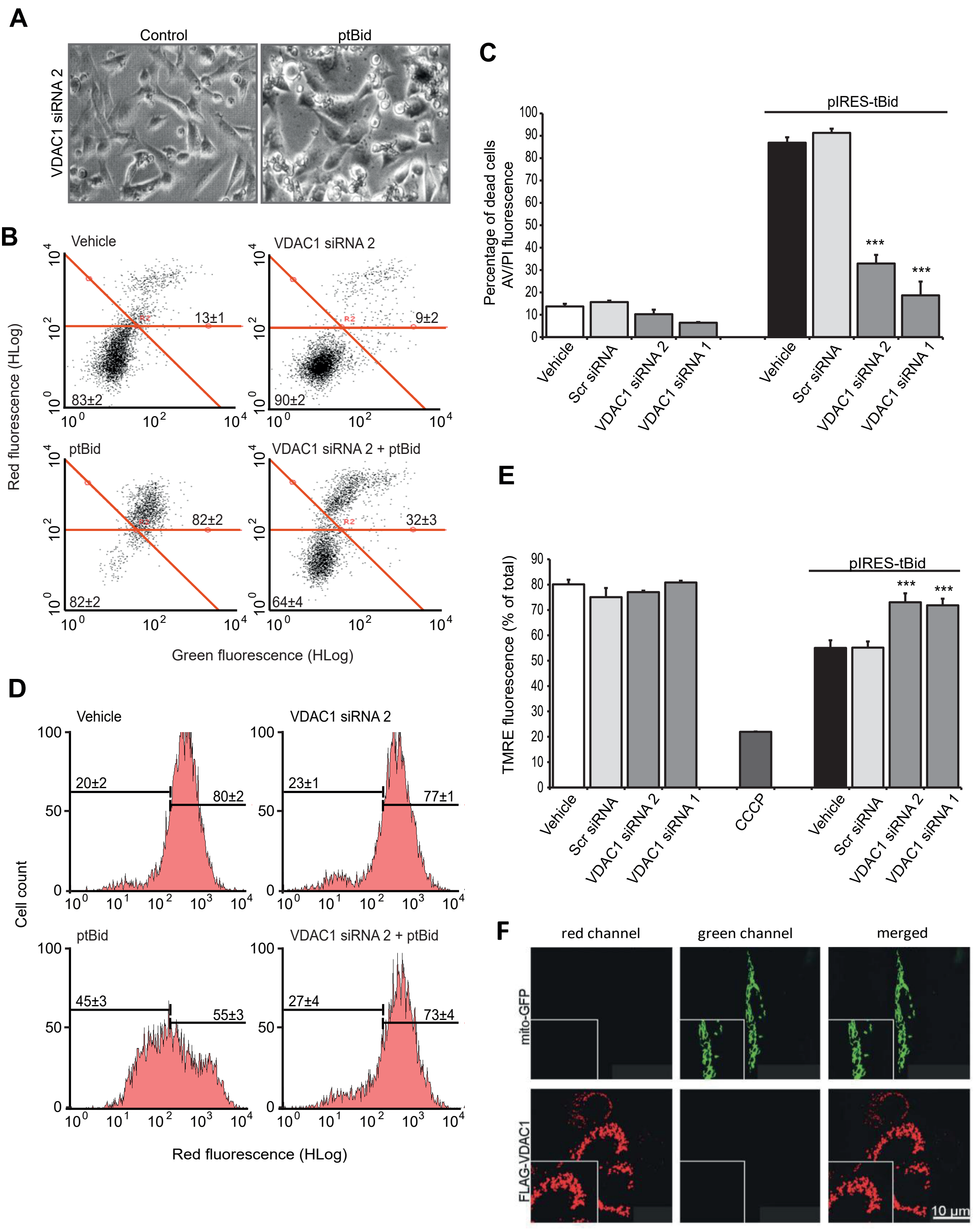
A
Control
ptBid
VDAC1 siRNA 2
B
Vehicle
VDAC1 siRNA 2
ptBid
VDAC1 siRNA 2 + ptBid
13±1
83±2
9±2
90±2
82±2
82±2
32±3
64±4
Red fluorescence (HLog)
Green fluorescence (HLog)
C
pIRES-tBid
Percentage of dead cells AV/PI fluorescence
***
***
Vehicle
Scr siRNA
VDAC1 siRNA 2
VDAC1 siRNA 1
D
Vehicle
VDAC1 siRNA 2
ptBid
VDAC1 siRNA 2 + ptBid
20±2
80±2
23±1
77±1
45±3
55±3
27±4
73±4
Cell count
Red fluorescence (HLog)
E
pIRES-tBid
TMRE fluorescence (% of total)
CCCP
F
red channel
green channel
merged
mito-GFP
FLAG-VDAC1
10 µm

### Figure S4

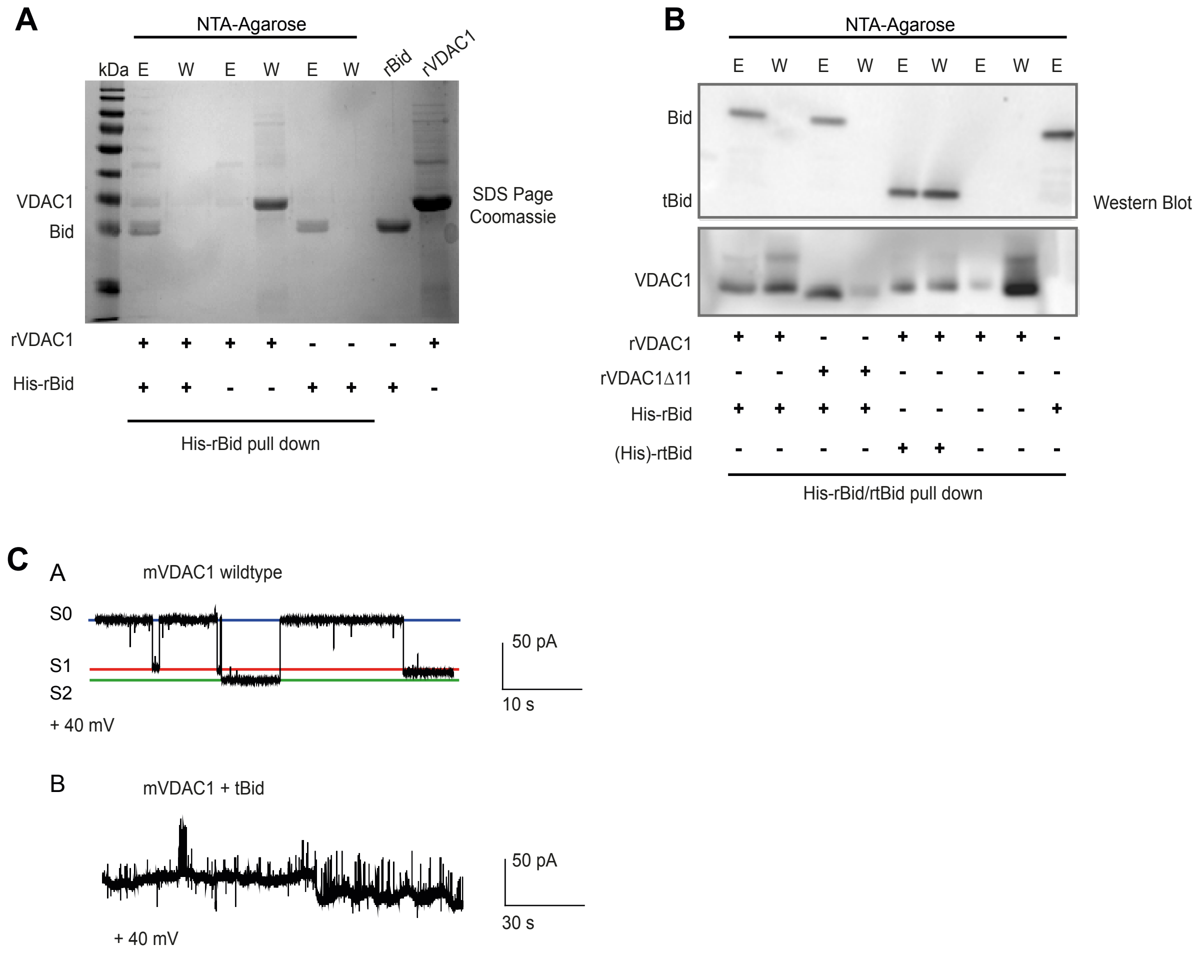

A
NTA-Agarose
kDa
E
W
E
W
E
W
rBid
rVDAC1
VDAC1
Bid
SDS Page
Coomassie
rVDAC1
+ + + + - - - +
His-rBid
+ + - - + + + -
His-rBid pull down
B
NTA-Agarose
E W E W E W E W E
Bid
tBid
Western Blot
VDAC1
rVDAC1
+ + - - + + + + -
rVDAC1Δ11
- - + + - - - - -
His-rBid
+ + + + - - - - +
(His)-rtBid
- - - - + + - - -
His-rBid/rtBid pull down
C
A
mVDAC1 wildtype
S0
S1
S2
+ 40 mV
50 pA
10 s
B
mVDAC1 + tBid
50 pA
30 s
+ 40 mV

**Figure S5**

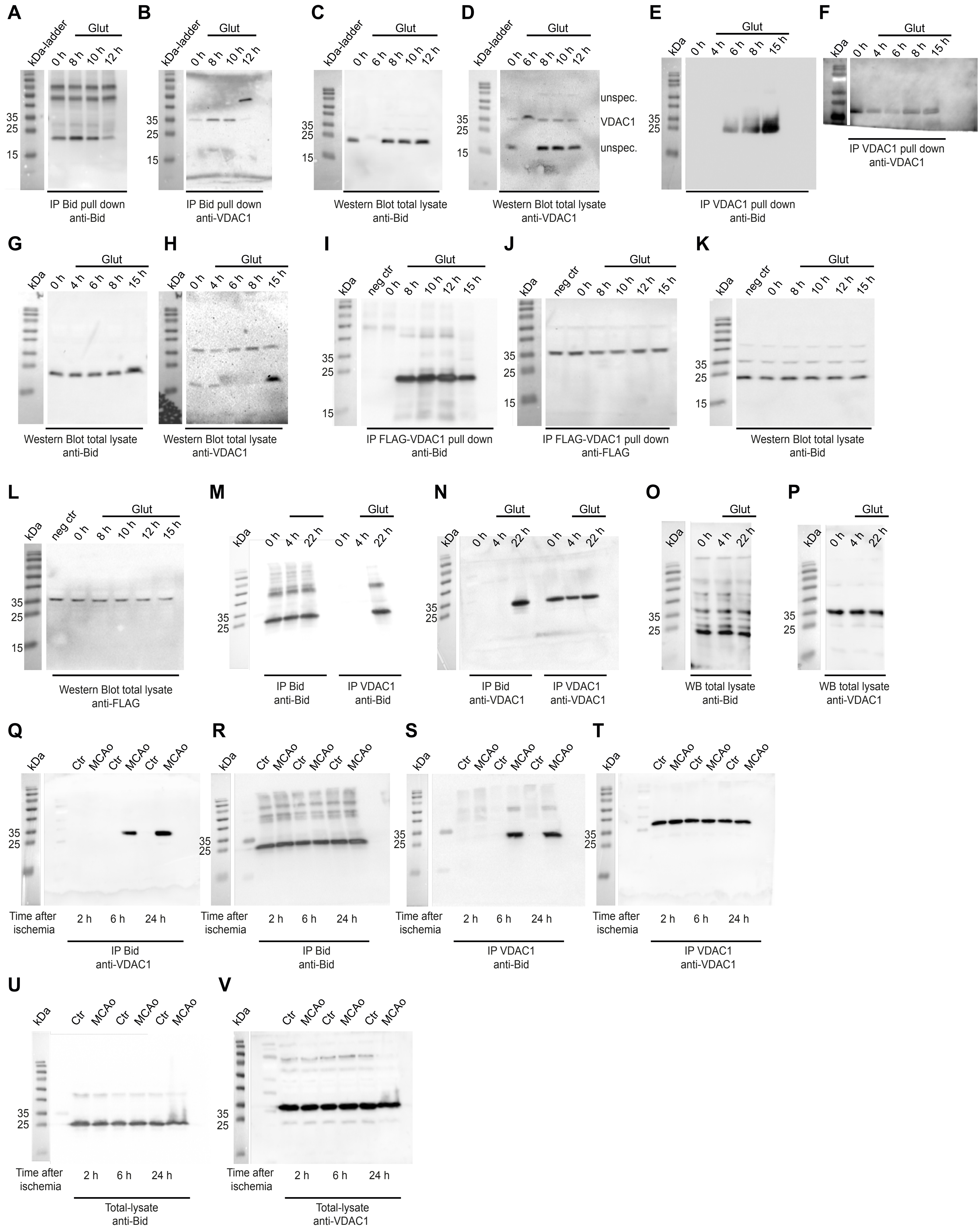
A
kDa-ladder
Glut
0 h
8 h
10 h
12 h
35
25
15
IP Bid pull down
anti-Bid
B
kDa-ladder
Glut
0 h
8 h
10 h
12 h
35
25
15
IP Bid pull down
anti-VDAC1
C
kDa-ladder
Glut
0 h
6 h
8 h
10 h
12 h
35
25
15
Western Blot total lysate
anti-Bid
D
kDa-ladder
Glut
0 h
6 h
8 h
10 h
12 h
35
25
15
unspec.
VDAC1
unspec.
Western Blot total lysate
anti-VDAC1
E
kDa
Glut
0 h
4 h
6 h
8 h
15 h
35
25
IP VDAC1 pull down
anti-Bid
F
kDa
Glut
0 h
4 h
6 h
8 h
15 h
IP VDAC1 pull down
anti-VDAC1
G
kDa
Glut
0 h
4 h
6 h
8 h
15 h
Western Blot total lysate
anti-Bid
H
kDa
Glut
0 h
4 h
6 h
8 h
15 h
Western Blot total lysate
anti-VDAC1
I
kDa
neg ctr
Glut
0 h
8 h
10 h
12 h
15 h
35
25
15
IP FLAG-VDAC1 pull down
anti-Bid
J
kDa
neg ctr
Glut
0 h
8 h
10 h
12 h
15 h
35
25
15
IP FLAG-VDAC1 pull down
anti-FLAG
K
kDa
neg ctr
Glut
0 h
8 h
10 h
12 h
15 h
35
25
15
Western Blot total lysate
anti-Bid
L
kDa
neg ctr
Glut
0 h
8 h
10 h
12 h
15 h
35
25
15
Western Blot total lysate
anti-FLAG
M
kDa
Glut
0 h
4 h
22 h
0 h
4 h
22 h
35
25
IP Bid
anti-Bid
IP VDAC1
anti-Bid
N
kDa
Glut
Glut
0 h
4 h
22 h
0 h
4 h
22 h
35
25
IP Bid
anti-VDAC1
IP VDAC1
anti-VDAC1
O
kDa
Glut
0 h
4 h
22 h
35
25
WB total lysate
anti-Bid
P
kDa
Glut
0 h
4 h
22 h
35
25
WB total lysate
anti-VDAC1
Q
kDa
Ctr
MCAo
Ctr
MCAo
Ctr
MCAo
35
25
Time after ischemia
2 h
6 h
24 h
IP Bid
anti-VDAC1
R
kDa
Ctr
MCAo
Ctr
MCAo
Ctr
MCAo
35
25
Time after ischemia
2 h
6 h
24 h
IP Bid
anti-Bid
S
kDa
Ctr
MCAo
Ctr
MCAo
Ctr
MCAo
35
25
Time after ischemia
2 h
6 h
24 h
IP VDAC1
anti-Bid
T
kDa
Ctr
MCAo
Ctr
MCAo
Ctr
MCAo
35
25
Time after ischemia
2 h
6 h
24 h
IP VDAC1
anti-VDAC1
U
kDa
Ctr
MCAo
Ctr
MCAo
Ctr
MCAo
35
25
Time after ischemia
2 h
6 h
24 h
Total-lysate
anti-Bid
V
kDa
Ctr
MCAo
Ctr
MCAo
Ctr
MCAo
35
25
Time after ischemia
2 h
6 h
24 h
Total-lysate
anti-VDAC1
